# Comprehensive Benchmarking and Ensemble Approaches for Metagenomic Classifiers

**DOI:** 10.1101/156919

**Authors:** Alexa B. R. McIntyre, Rachid Ounit, Ebrahim Afshinnekoo, Robert J. Prill, Elizabeth Hénaff, Noah Alexander, Sam Minot, David Danko, Jonathan Foox, Sofia Ahsanuddin, Scott Tighe, Nur A. Hasan, Poorani Subramanian, Kelly Moffat, Shawn Levy, Stefano Lonardi, Nick Greenfield, Rita R. Colwell, Gail L. Rosen, Christopher E. Mason

## Abstract

**Background:** One of the main challenges in metagenomics is the identification of microorganisms in clinical and environmental samples. While an extensive and heterogeneous set of computational tools is available to classify microorganisms using whole genome shotgun sequencing data, comprehensive comparisons of these methods are limited. In this study, we use the largest (n=35) to date set of laboratory-generated and simulated controls across 846 species to evaluate the performance of eleven metagenomics classifiers. We also assess the effects of filtering and combining tools to reduce the number of false positives.

**Results:** Tools were characterized on the basis of their ability to (1) identify taxa at the genus, species, and strain levels, (2) quantify relative abundance measures of taxa, and (3) classify individual reads to the species level. Strikingly, the number of species identified by the eleven tools can differ by over three orders of magnitude on the same datasets. However, various strategies can ameliorate taxonomic misclassification, including abundance filtering, ensemble approaches, and tool intersection. Indeed, leveraging tools with different heuristics is beneficial for improved precision. Nevertheless, these strategies were often insufficient to completely eliminate false positives from environmental samples, which are especially important where they concern medically relevant species and where customized tools may be required.

**Conclusions:** The results of this study provide positive controls, titrated standards, and a guide for selecting tools for metagenomic analyses by comparing ranges of precision and recall. We show that proper experimental design and analysis parameters, including depth of sequencing, choice of classifier or classifiers, database size, and filtering, can reduce false positives, provide greater resolution of species in complex metagenomic samples, and improve the interpretation of results.

## Background

Sequencing has helped researchers identify microorganisms with roles in such diverse areas as human health [1], the color of lakes [2], and climate [3,4]. The main objectives when sequencing a metagenomic community are to accurately detect, identify, and describe its component taxa fully and accurately. False positives, false negatives, and speed of analysis are critical concerns, in particular when sequencing is applied to medical diagnosis or tracking infectious agents.

Selective amplification (e.g. 16S, 18S, ITS) of specific gene regions has long been standard for microbial community sequencing, but it introduces bias and omits organisms and functional elements from analysis. Recent large-scale efforts to characterize the human microbiome [5] and a variety of Earth microbiomes [6] used the 16S genes of ribosomal RNA (rRNA) as amplicons. Highly conserved regions within these genes permit the use of common primers for sequencing [7]. Yet certain species of archaea include introns with repetitive regions that interfere with the binding of the most common 16S primers [8,9] and 16S amplification is unable to capture viral, plasmid, and eukaryotic members of a microbial community, which may represent pivotal drivers of an individual infection or epidemic [10]. 16S amplification is also often insufficient for discrimination at the species and strain levels of classification. Although among closely-related strains of prokaryotes, conserved genes with higher evolutionary rates than 16S rRNA [11] or gene panels could improve discriminatory power, these strategies suffer from low adoption and underdeveloped reference databases.

While whole genome shotgun sequencing addresses some of the issues associated with amplicon-based methods, other challenges arise. Amplification-based methods remain a cheaper option, and 16S databases are more extensive than shotgun databases. Also, taxonomic annotation of short reads produced by most standard sequencing platforms remains problematic, since shorter reads are more likely to map ambiguously to related taxa that are not actually present in a sample. Strategies for whole genome shotgun classification rely on several methods, including alignment (to all sequences or taxonomically unique markers), composition (*k*-mer analysis), phylogenetics (using models of sequence evolution), assembly, or a combination of these methods. Analysis tools focusing on estimation of abundance tend to use marker genes, which decreases the number of reads classified but increases speed [12]. Tools that classify at the read-level have applications beyond taxonomic identification and abundance estimation, such as identifying contaminating reads for removal before genome assembly, calculating coverage, or determining the position of bacterial artificial chromosome clones within chromosomes [13,14].

Environmental surveys of the New York City (NYC) subway system microbiome and airborne microbes found that metagenomic analysis tools unable to find a match to any reference genome for about half of input reads to any reference genome, demonstrating the complexity of the data and limitations of current methods and databases [15,16]. Environmental studies also highlight the importance of reliable species identification when determining pathogenicity. All analysis tools used in the initial NYC subway study detected matches to sequences or markers associated with human pathogens in multiple samples, although subsequent analyses by the original investigators, as well as others, showed there was greater evidence for related, but non-pathogenic, organisms [17–19]. The problem of false positives in metagenomics has been recognized and reported [20,21]. Strategies including filtering and combining classifiers have been proposed to correct the problem but a thorough comparison of these strategies has not been done. Recent publications have focused on detecting and identifying harmful or rare microorganisms [19,21,22]. However, when studying common non-pathogenic microbes, investigators routinely rely on the accuracy of increasingly rapid analyses from metagenomic classifiers [22].

Efforts to standardize protocols for metagenomics, including sample collection, nucleic acid extraction, library preparation, sequencing, and computational analysis are underway, including large-scale efforts like the Microbiome Quality Control (MBQC) [23], the Genome Reference Consortium (GRC), the International Metagenomics and Microbiome Standards Alliance (IMMSA) [24], the Critical Assessment of Metagenomics Interpretation (CAMI), and others [2,23,25–27]. Comparisons of available bioinformatics tools have only recently been published [20,12,27–29]. For example, Lindgreen, et al. (2016) evaluated a set of fourteen metagenomics tools, using six datasets comprising more than 400 genera, with the analysis limited to phylum and genus. A similar study done by Peabody et al. (2015) evaluated algorithms to the species level but included only two data sets representing eleven species, without taking into account the evolution of the taxonomy of those species [30]. Meanwhile, the number of published tools for the identification of microorganisms continues to increase. At least eighty tools are currently available [27], although some are no longer maintained. Publications describing new methods tend to include comparisons to only a small subset of existing tools, ensuring an enduring challenge of determining which tools should be considered “state-of-the-art” for metagenomics analysis.

To address the challenge, we curated and created a set of fourteen laboratory-generated and twenty-one simulated metagenomic standards datasets, comprising 846 species, including read- and strain-level annotations for a subset of datasets and sequences for a new, commercially available DNA standard that includes bacteria and fungi (Zymo BioOMICs). We further tested tool agreement using a deeply-sequenced (>100M reads) environmental sample, and developed new ensemble “voting” methods for improved classification. These data provide an online resource for extant tools and are freely available (http://ftp-private.ncbi.nlm.nih.gov/nist-immsa/IMMSA/) for others to use for benchmarking future tools or new versions of current tools.

## Results

We compared the characteristics and parameters of a comprehensive set of eleven metagenomic tools [13,32–43] (Table 1) representing a variety of classification approaches (*k*-mer composition, alignment, marker). We also present a comprehensive evaluation of their performance, using both simulated and biological datasets, across a wide range of GC content, complexity, and species similarity characteristics (Supplementary Table 1).

**Table 1.**
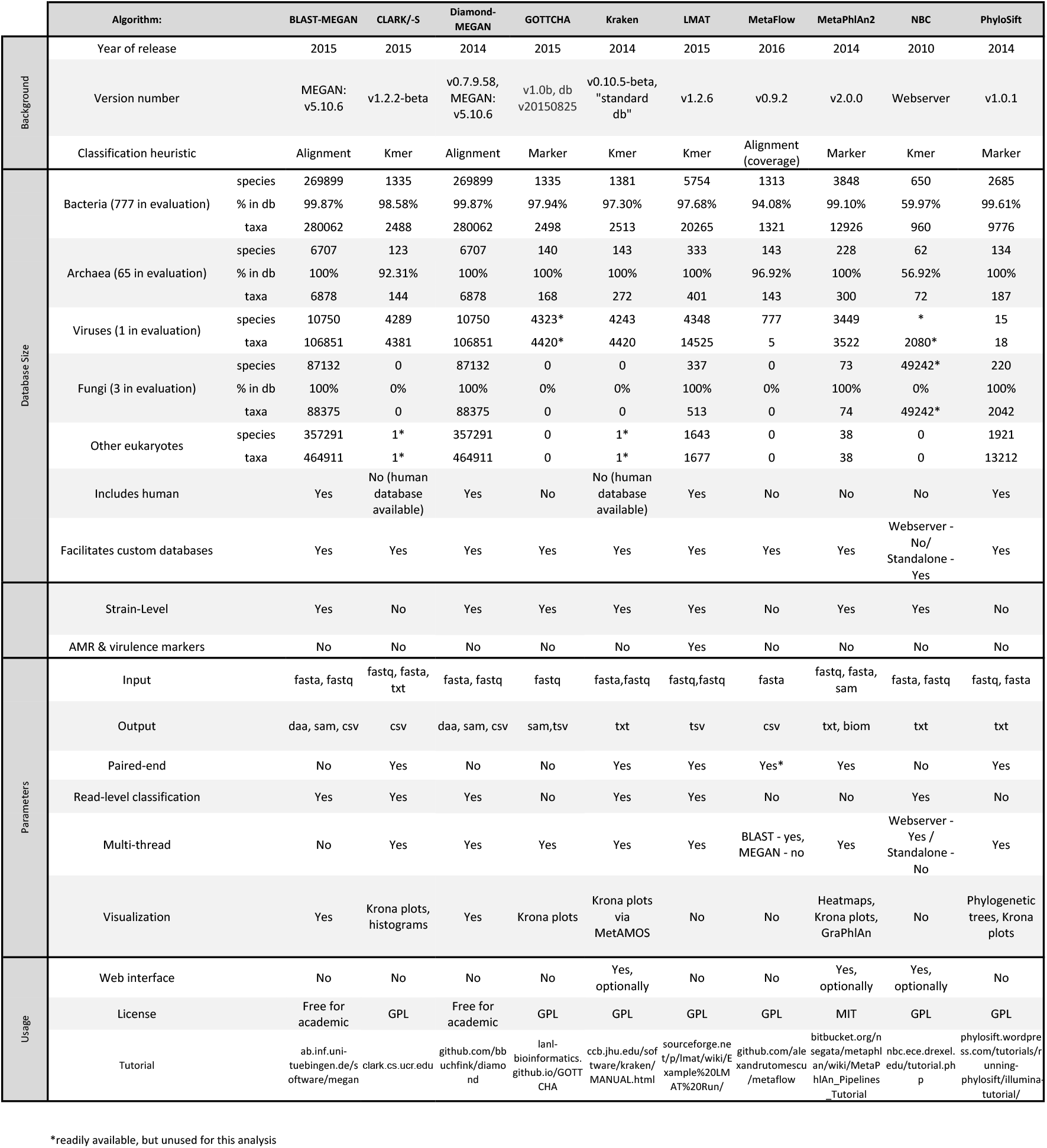
Comparison of tools by classification strategies and associated databases.

### Genus, Species, and Subspecies Level Comparisons

From the platypus [21] to *Yersinia pestis* [16], false positives can plague metagenomic analyses. To evaluate the extent of the problem with respect to specific tools, we calculated precision and recall based on detection of the presence or absence of a given genus, species, or subspecies, first if detected at any abundance. We calculated precision, recall, area under the precision-recall curve (AUPR), and F1 score for each tool across all 35 datasets for all 846 species. When compared by mean AUPR (mAUPR), all tools performed best at the genus level (45.1% ≤ mAUPR ≤ 86.6%, Figure 1a), with small decreases in performance at the species level (40.1% ≤ mAUPR ≤ 84.1%, Figure 1b). Calls at the subspecies (strain) level showed a more marked decrease on all measures for the subset of datasets that included complete strain information (17.3% ≤ mAUPR ≤ 62.5%, Figure 1c). For *k*-mer-based tools, adding an abundance threshold improved precision and increased the F1 score, which is more affected than AUPR by false positives detected at low abundance, bringing them to the same range as marker-based tools, which tended to be more precise (Figure 1d-e).

**Figure 1.**
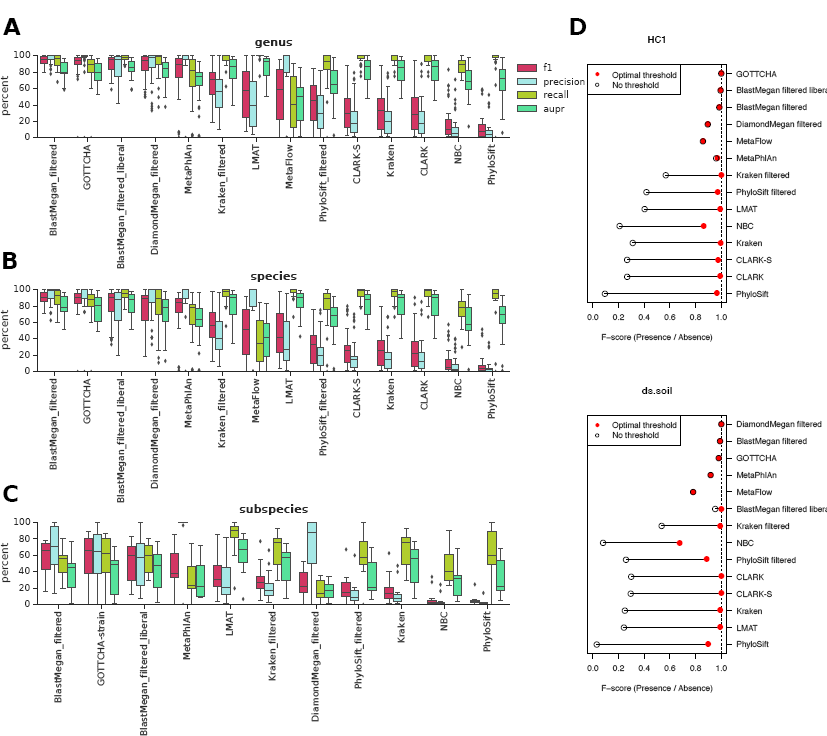
The F1-score, precision, recall, and area under the precision-recall curve (where tools are sorted by decreasing mean F1-score) across datasets with available truth sets for taxonomic classifications at the **a)** genus (35 datasets), **b)** species (35 datasets), and **c)** subspecies (12 datasets) levels. **d)** The F1 score changes depending on relative abundance thresholding, as shown for two datasets. The upper bound in red marks the optimal abundance threshold to maximize F1 score, adjusted for each dataset and tool. The lower bound in black indicates the F1 score for the output without any threshold. Results are sorted by the distance between upper and lower bounds.

### Performance Across Datasets

Grouping datasets into simulated reads and biological samples revealed that precision is notably lower for biological samples that are titrated and then sequenced (Supplementary Figure 1). We initially hypothesized that tools would attain lower precision with biological data because (1) they detect true contaminants, (2) they detect close variants of the reference strain, or (3) simulated data does not fully capture errors, GC content range, and read distribution biases present in biological data. However, whether data was simulated had no significant effect on the number of false positives detected (Figure 2, with the exception of MetaFlow, which showed a significant trend only with outliers, Supplementary Figure 2a). The decrease in precision could instead occur because the biological samples contained fewer species on average, but tools detected similar numbers of false positives (no significant relationship was found between the number of taxa in a sample and false positives for most tools). False positives for almost all *k*-mer-based methods tended to increase with more reads (e.g., Supplementary Figure 2b), confirming the relationship between depth and total number of misclassified reads. Interestingly, the same relationship does not exist for most marker- and alignment-based read callers, suggesting any additional reads that are miscalled are miscalled as the same species as read depth increases. BLAST-MEGAN and PhyloSift without or with laxer filters were exceptions, but filtering was sufficient to avoid the trend. On further examination, the significant relationship between number of taxa and read length and false positive counts for MetaPhlAn and GOTTCHA appeared weak for MetaPhlAn and entirely due to outliers for GOTTCHA (Supplementary Figure 2c-f), indicating such misclassification can be very dataset-specific.

**Figure 2.**
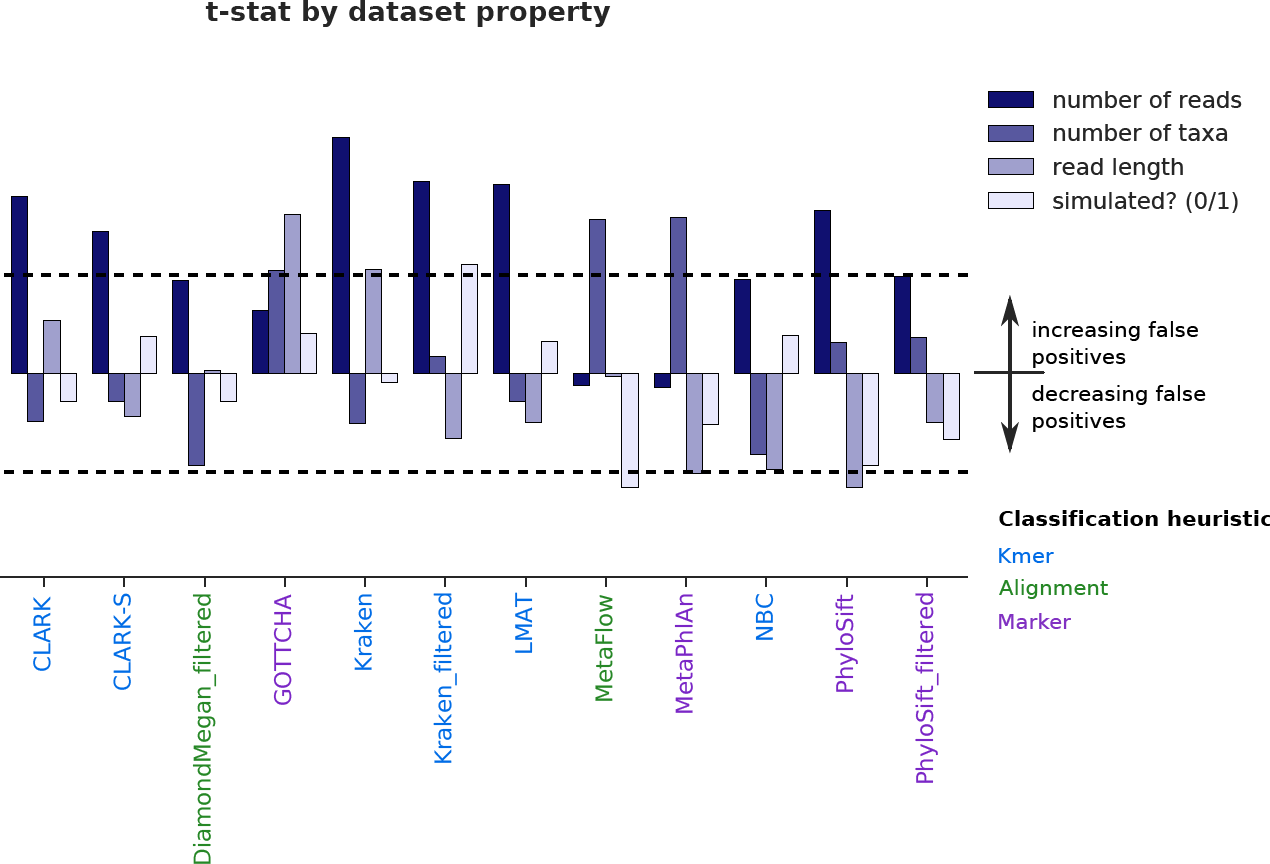
Number of false positives called by different tools as a function of dataset features. The t-statistic for each feature is reported after fitting a negative binomial model, with p-value > 0.05 within the dashed lines and significant results beyond.

The mAUPR for each sample illustrates wide variation among datasets (Supplementary Tables 1 and 2, Supplementary Figure 3). Difficulty in identifying taxa was not directly proportional to number of species in the sample, as evidenced by the fact that biological samples containing ten species and simulated datasets containing 25 species with lognormal distributions of abundance were among the most challenging (lowest mAUPR). Indeed, some datasets had a rapid decline in precision for almost all tools (e.g. LC5), which illustrates the challenge of calling species with low depth coverage and the potential for improvement using combined or ensemble methods.

### Ensemble Approaches to Determine Number and Identity of Species Present

To gauge the benefits of combining multiple tools for accuracy and measuring the actual number of species present in a sample, a series of tests was used. First, a combination of five lower-precision tools (CLARK, LMAT, PhyloSift, Kraken, and NBC) showed the overlap between species in the truth sets. These combined results showed that the most abundant species identified by the tools was relatively high for subset sizes close to the actual number of species (Figure 3a). Overlap in the number of species identified by all five tool 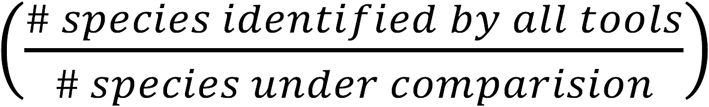 was evaluated by sorting species according to abundance and varying the number of results included in the comparison (Figure 3b). For most samples, discrepancies in results between tools were higher and inconsistent below the known number of species because of differences in abundance estimates. Discrepancies also increased steadily as evaluation size exceeded the actual number of species to encompass more false positives. Thus, these data show that the rightmost peak in percent overlap in even lower-precision tools approximates the known, true number of species (Figure 3c). However, the number of species identified by high-precision tools provides a comparable estimate of species diversity and number, with GOTTCHA and filtered results for BLAST-MEGAN and Diamond-MEGAN still outperforming the combined-tool strategy (Figure 3d).

**Figure 3.**
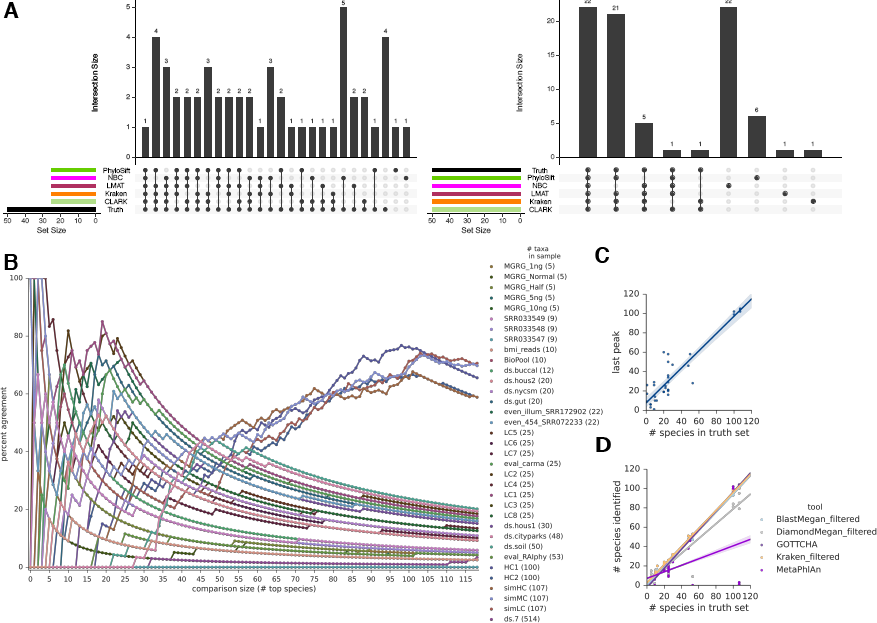
Combining results from imprecise tools can predict the true number of species in a dataset. **a)** UpSet plots of the top-X (by abundance) species uniquely found by a classifier or group of classifiers (grouped by black dots at bottom, unique overlap sizes in the bar charts above). The eval_RAIphy dataset is presented as an example, with comparison sizes X=25 and X=50. The percent overlap, calculated as the number of species overlapping between all tools, divided by the number of species in the comparison, increases around the number of species in the sample (50 in this case). **b)** The percent overlaps for all datasets show a similar trend. **c)** The rightmost peak in **b** approximates the number of species in a sample, with a root mean square error of 8.9 on the test datasets. **d)** Precise tools can offer comparable or better estimates of species count. Root mean square error = 3.2, 3.8, 3.9, 12.2, 32.9 for Kraken filtered, BlastMegan filtered, GOTTCHA, DiamondMegan filtered, and MetaPhlAn2, respectively.

Additional pairwise combinations of tools also show general improvements in taxonomic classification, with the overlap between pairs of tools almost always increasing precision compared to results from individual tools (Figure 4a). At the species level, combining filtered BLAST-MEGAN and Diamond-MEGAN, NBC, or GOTTCHA, and GOTTCHA or Diamond-MEGAN increased mean precision to over 95%, while thirty other combinations increased precision to over 90%. However, depending on the choice of tools, improvement in precision was incremental at best. For example, combining two *k*-mer-based methods (e.g., CLARK-*S* and NBC, with mean precision 26.5%) did not improve precision to the level of most of the marker-based tools. Increases in precision were offset by decreases in recall, notably when tools with small databases such as NBC were added (Figure 4b). Overall, this shows that leveraging tools with different heuristics is beneficial for precision.

**Figure 4.**
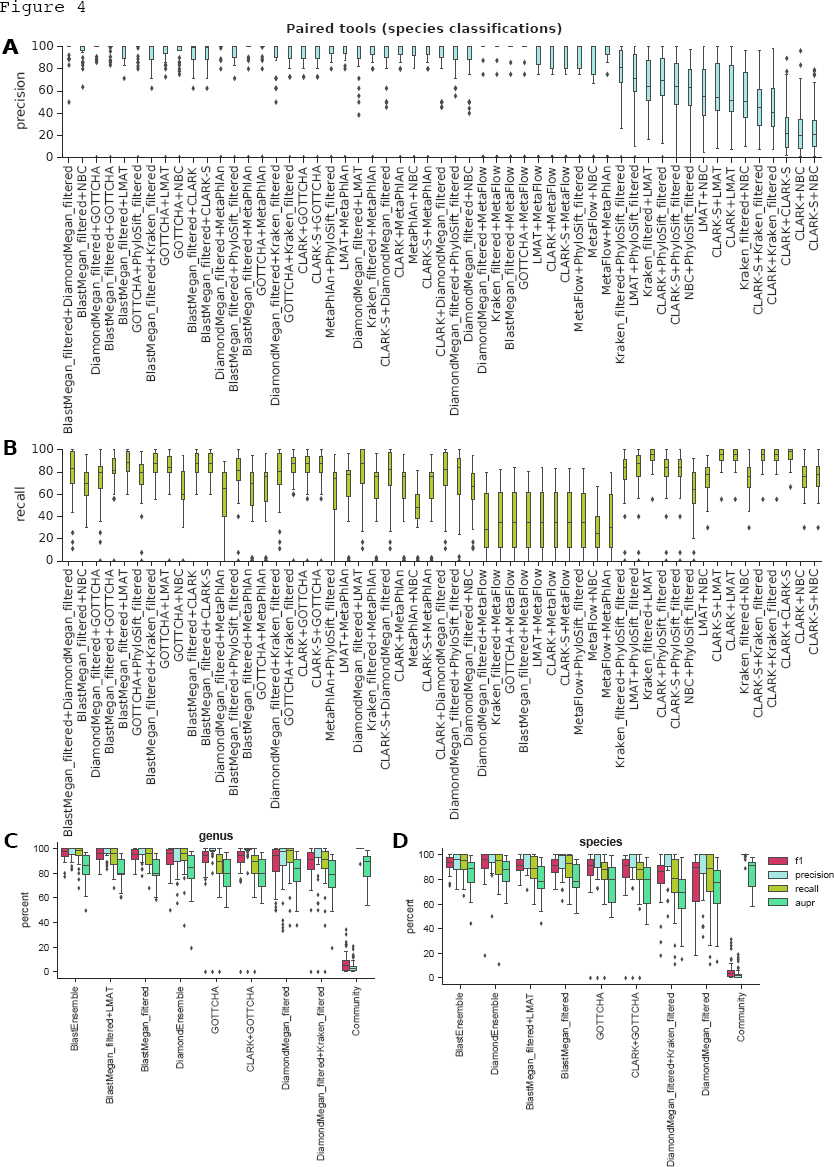
The **a)** precision and **b)** recall for intersections of pairs of tools at the species level, sorted by decreasing mean precision. A comparison between multitool strategies and combinations at the **(c)** genus and **(d)** species levels. The top unique (non-overlapping) pairs of tools by F1 score from **(a-b)** are benchmarked against the top single tools at the species level by F1 score, ensemble classifiers that take the consensus of four or five tools (see *Methods),* and a community predictor that incorporates the results from all eleven tools in the analysis to improve AUPR.

We next designed a community predictor that combines abundance rankings across all tools (see *Methods, Scripts).* Consensus ranking offered improvement over individual tools in terms of mAUPR, which gives an idea of the accuracy of abundance rankings (Supplementary Table 2). Unlike pairing tools, this approach can also compensate for variations in database completeness among tools for samples of unknown composition, since detection at high abundance by only a subset of tools was sufficient for inclusion in the filtered results of the community predictor. However, by including every species called by any tool, precision inevitably falls.

As alternatives, we designed two “majority vote” ensemble classifiers using the top tools by F1 score either including BLAST (the slowest tool) or not. At the genus-level (Figure 4c), the majority vote BlastEnsemble had the best F1 score due to limited loss in precision with improved recall. However, we show that little performance is sacrificed using only BLAST-MEGAN or the overlap of BLAST-MEGAN and LMAT. If one does not want to run BLAST for speed reasons, the majority vote DiamondEnsemble is a competitive alternative, improving the F1 score over Diamond-MEGAN or GOTTCHA alone. At the species-level (Figure 4d), the Blast Ensemble and Diamond Ensemble ranked highest, showing that BLAST can be avoided while maintaining comparable performance. Finally, pairing tools could occasionally lead to worse performance; for example, GOTTCHA combined with CLARK gave lower F1 score than just GOTTCHA (Figure 4d).

### Classifier Performance by Taxa

We next sought to discern the consistently hardest species to detect within and across the tools; the performance of each classifier by taxa are provided in the supplementary material. The most difficult taxa to identify at each taxonomic level (averaged over all classifiers) are *Archaea* (Superkingdom), *Acidobacteria* (phylum), *Acidobacteriia* (class), *Acidobacteriales* (order), *Crocosphaera* (genus), and *Acinetobacter sp. NCTC 10304/Corynebacterium pseudogenitalium/Propionibacterium sp. 434-HC2* (species). Common taxa such as Proteobacteria/Firmicutes/ Actinobacteria (phylum) and *Lactobacillus/Staphylococcus/Streptococcus* (species) were frequent false positives. Classifiers show bias towards these taxa likely because they are better represented in databases than others. In terms of false negatives, it is interesting to note that genera that include highly similar species, such as *Bacillus, Bifidobacterium,* and *Shigella* were commonly miscalled.

### Negative Controls

Of the methods tested, seven did not include the human genome in their default database. For those that did, human DNA was identified as the most abundant species in the biological negative controls, sequenced using a human reference material (NA12878) spiked into a MoBio PowerSoil extraction kit (Table 2). Most of the tools identified additional non-human species, between a mean of 4.67 for GOTTCHA and 1,360 for CLARK-S. MetaFlow and BLAST-MEGAN (default filter) provided the only results that did not identify additional species. Notably, not all additional species are necessarily false positives; previous studies (e.g., [44]) detected biological contaminants in sequencing data.

**Table 2.**
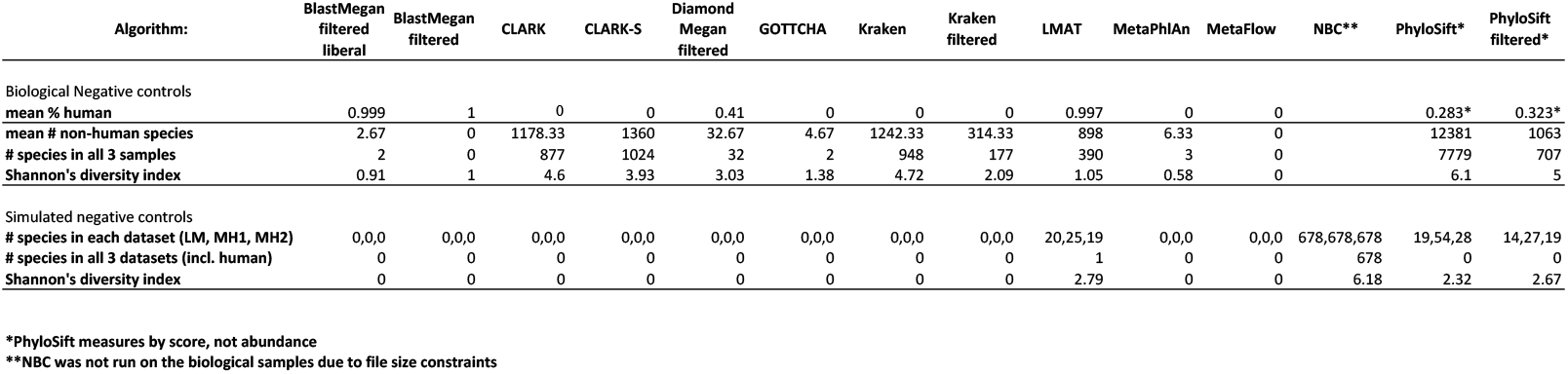
Negative control results on biological samples with human DNA spiked in and simulated data constructed from nullomers (17-mers that do not map to any reference).

Using pairs of tools with mean precision greater than 95% (n = 17) on the test datasets at the genus level, we found *Acinetobacter* and *Escherichia* were genera of putative sequencing and/or reagent contaminants. Previous studies have also detected contamination with both [44]. High-precision pairs at the species level (n = 6) reported only *Escherichia coli.*

We next tested a set of three million simulated negative control sequences that do exist in any known species (see *Methods*, Table 2). Most tools did not identify any species in these synthetic control sequences, although PhyloSift, NBC, and LMAT identified false positives at low probability scores (PhyloSift) or abundances (NBC and LMAT). The identification of *Sorangium cellulosum* as the most abundant species in all three datasets indicates size bias among NBC’s false positives. The *S. cellulosum* genome is particularly large for bacteria at 13M base pairs [45]. Further top ranking species from NBC were consistent despite smaller genomes than other organisms in the database, most likely because there are more reference sequences available at the subspecies level for these common microbes (29 *E. coli,* and 9 *B. cereus* in the NBC database). LMAT consistently identified human as the most abundant species in all three datasets without any other overlap between the datasets, suggesting a bias towards the host reference genome. PhyloSift results were variable, with no species consistently reported in all three datasets.

### Detection of Pathogenic False Positives

We next applied similar methods of combining tool predictions in an attempt to rule out false positives in two datasets in which *Ba cillus anthracis* had previously been reported by BLAST, MetaPhlAn, and SURPI [16]. Interestingly, all eleven tested tools also detected the pathogenic taxa *Bacillus anthracis* or *Bacillus cereus biovar anthracis* in at least one of the datasets without filtering, although the number of reads and relative abundance were low for most tools (Supplementary Table 3). Moreover, the tools still reported the organisms after applying tool-suggested filters. Even taking pairs of tools with high precision for known samples showed that two of six such pairs still reported anthrax, while still more reported either *B. anthracis* or *B. cereus biovar anthracis,* further underscoring the need for caution even when many tools identify pathogenic organisms in a sample. Inference based on plasmids and specific genetic markers can more accurately separate harmless from pathogenic strains, even within the monophyletic species of the genus *Bacillus.* To address this challenge, we have released a new tool that can accurately discriminate harmless from pathogenic strains of *Bacillus* using plasmid and specific gene markers [19].

### Relative Abundance

After defining the parameters for species detection and counting, we calculated the accuracy of relative abundance predictions (Figure 5a-b) for titrated and simulated samples. Almost all tools could predict the percentage of a species in a sample within a few percentage points, yet with higher variance of the predicted value at low abundance. GOTTCHA was an exception, performing poorly with log-normally distributed samples (Figure 5a, c) despite success with more evenly distributed samples (Figure 5b). Although GOTTCHA showed promise in relative abundance estimation on first publication [29], our results are consistent with those from Lindgreen, et al. (2016) at higher levels of classification (phylum and genus). While the log-modulus examines a fold-change, the L1 distance shows the distance between relative abundance vectors by dataset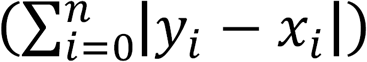, where *y* is the expected profile and x observed profile (Figure 5d) [46]. Many tools showed greater variation between datasets, as measured by the L1 distance for simulated datasets, especially BLAST and Diamond. The ensemble methods performed the best on the simulated data but had more variation than NBC, MetaPhlAn, and CLARK. On the biological samples, DiamondEnsemble was competitive but again had greater deviation than CLARK and tended to underestimate the relative abundance while CLARK tended to overestimate.

**Figure 5.**
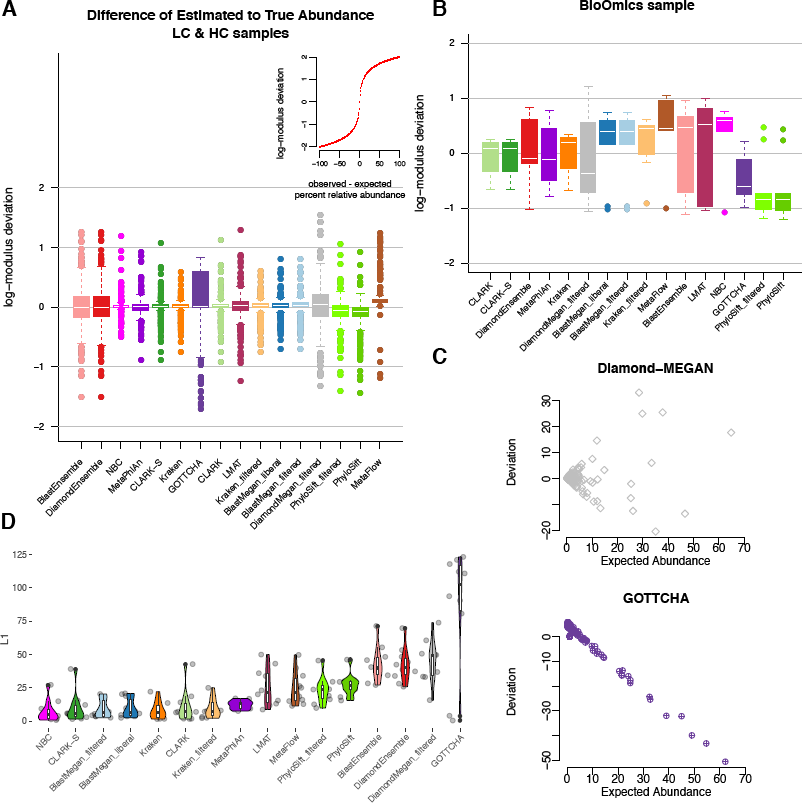
The relative abundances of species detected by tools compared to their known abundances for **a)** simulated datasets and **b)** a biological dataset, sorted by median log-modulus difference (difference’ = sign(difference)*log(1+|difference|)). Most differences between observed and expected abundances fell between 0 and 10, with a few exceptions (see inset for scale). **c)** The deviation between observed and expected abundance by expected percent relative abundance for two high variance tools on the simulated data. While most tools, like Diamond-MEGAN, did not show a pattern in errors, GOTTCHA overestimated low-abundance species and underestimated high-abundance species in the log-normally distributed data. **d)** The L1 distances between observed and expected abundances show the consistency of different tools across simulated datasets.

### Limits of Detection and Depth of Sequencing

To quantify the amount of input sequence required for detection, recall was calculated as a function of sequencing depth for each input organism, using the Huttenhower HC/LC datasets (Figure 6a). Each bin represents 17-69 input organisms, for a total of 197 organisms in the analysis. In general, *k*-mer-based methods (CLARK, Kraken, and LMAT) produced the highest recall, while other methods required higher sequencing depth to achieve equivalent recall.

**Figure 6.**
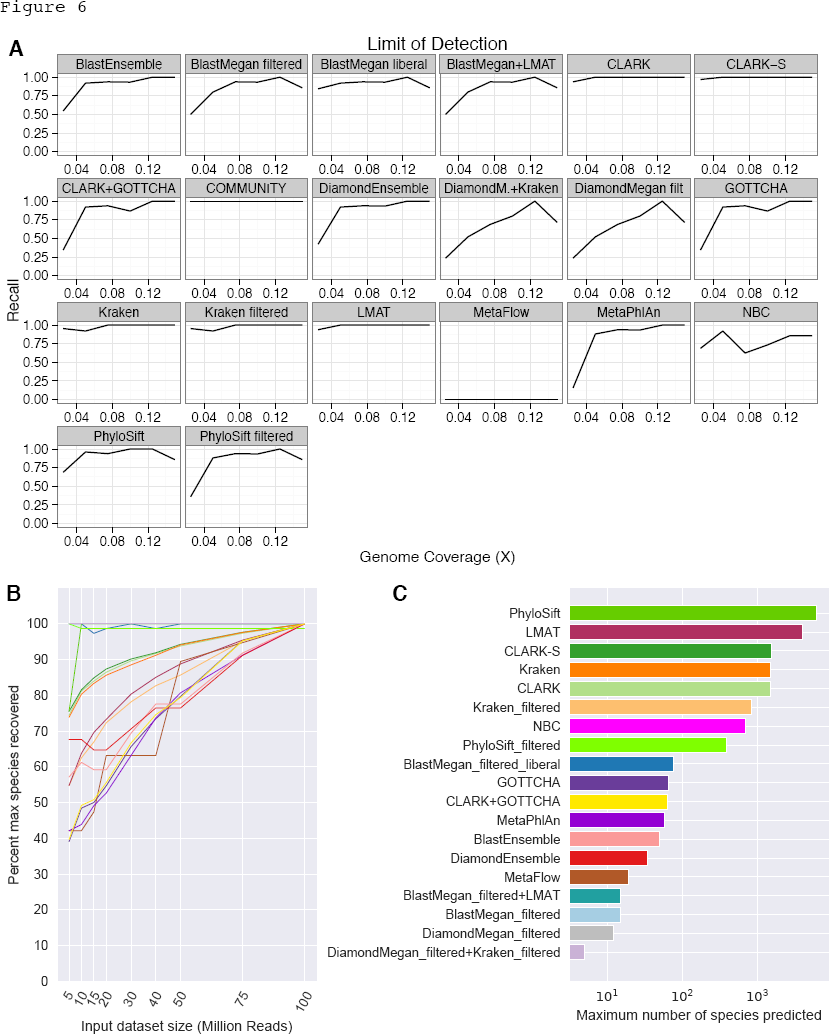
**a)** Recall at varying levels of genome coverage on the HC and LC datasets (using the least filtered sets of results for each tool). **b)** Downsampling a highly sequenced environmental sample shows depth of sequencing significantly affects results for specific tools, expressed as a percentage of the maximum number of species detected. Depending on strategy, filters can decrease the changes with depth. **c)** The maximum number of species detected by each tool at any depth.

Yet, depth of sequencing can significantly change the results of a metagenomic study, depending on the tool used. Using a deeply-sequenced, complex environmental sample from the Gowanus Canal (100M reads), we subsampled the full dataset (to 5, 10, 15, 20, 30, 40, and 50M reads) to identify the depth at which each tool recovered its maximum number of predicted species (Figure 6b). Reinforcing our analysis of limits of detection, marker-based tools identified far more species as depth of sequencing increased, an effect slightly attenuated by filtering (Figure 6c). Among *k*-mer-based tools, LMAT showed the largest increase, while Kraken, CLARK, and CLARK-S showed more gradual increases. Filtering Kraken results decreased the absolute number of species identified but increased the slope of the trend. As investigators consider depth of sequencing in their studies, they should keep in mind that results will change, depending on the tool selected and method of filtering. Based on these results, standardizing sequencing depth is extraordinarily important to compare multiple samples within studies or from similar studies.

### Nanopore Reads

Short, highly accurate reads are the primary focus of most analysis tools but newer, long-read sequencing methods can offer a lower cost, more portable alternative for metagenomics studies [44]. We tested the tools using two titrated MGRG mixtures (five and eleven species, respectively) sequenced using one of the first available versions (R6) and a newer update (R9) of the MinION from Oxford Nanopore Technologies (Supplementary Figure 4). “2D” consensus-called reads from the initial release of the MinION attained around 80% alignment accuracy, increasing to around 95% since then. Most *k*-mer- and alignment-based tools identified all component species of the mixture at some level of abundance, although also reported false positives among the top five results. CLARK and Diamond-MEGAN performed as well with lower quality data, while other tools were not as robust. Classification of reads with an average quality score of >Q9 improved results for LMAT. Marker-based methods did not perform well, likely in part because the datasets were small and failed to cover the expected markers.

### Read-level Analysis

Finally, we used the output from eight tools that delineate results for each individual read to measure precision and recall for species identification at the read-by-read level, where 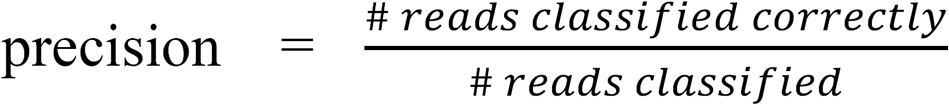 and 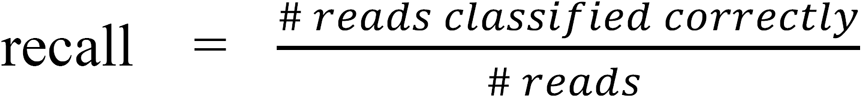 with classification to species or subspecies (Table 3). Both measures were high for all tools, although low recall was observed for some of the datasets, depending on whether the species in the dataset were also in a tool’s database. The low recall of some tools can also be explained by the low proportion of classified reads after filtering (e.g., Diamond-MEGAN and NBC). BLAST-MEGAN offered the highest precision, while CLARK-S most frequently provided highest recall. An ensemble approach was constructed by assigning each read to the most frequently called taxa among the different tools. Setting the quorum to one improved recall by 0.43% on average compared with results of best tool for each dataset, while maintaining precision comparable to the most precise tool for each dataset.

**Table 3.** Read-level analysis on twenty-one datasets for seven read-by-read classifiers: CLARK, CLARK-*S* (filtered), Kraken (filtered), LMAT (filtered), BLAST-MEGAN (filtered), Diamond-MEGAN (filtered), and NBC (filtered), as well as the ensemble classifiers (or meta-classifiers) that aim either to maximize precision or recall. Column 1 indicates the dataset name, column 2 shows the metric (either precision or recall), columns 3 to 9 provide the precision and recall for each dataset for all seven classifiers, column 10 and 11 record the precision and recall of the meta-classifier with the quorum value (in parentheses) that allows the highest precision or recall, respectively. Because shorter reads can ambiguously map to multiple species, optimal recall is 100 only for the unambiguously mapping datasets in which all reads should be mapped with high certainty to species.

| Dataset | Metric | CLARK | CLARK-S | Kraken | LMAT | BlastMegan | DiamondMegan | NBC | Meta-Classifer<br>(Precision) | Meta-Classifer<br>(Recall) |
| --- | --- | --- | --- | --- | --- | --- | --- | --- | --- | --- |
| HC1 | Precision | 99.73 | 97.79 | 99.93 | 99.70 | **99.98** | 97.94 | 94.83 | 99.41 (6) | 98.27 (1) |
|  | Recall | 85.10 | **90.30** | 74.16 | 74.57 | 77.38 | 23.92 | 62.42 | 88.91 (6) | 89.49 (1) |
| HC2 | Precision | 99.69 | 96.57 | 99.77 | 99.62 | **99.97** | 97.61 | 93.43 | 99.63 (6) | 97.02 (1) |
|  | Recall | 83.05 | **88.07** | 69.78 | 72.34 | 76.49 | 24.74 | 59.95 | 86.98 (6) | 87.46 (1) |
| LC1 | Precision | 95.42 | 94.23 | 94.36 | 95.84 | 95.39 | 97.55 | 94.75 | **99.93 (7)** | 98.93 (1) |
|  | Recall | 85.89 | **91.05** | 74.57 | 79.90 | 78.25 | 27.91 | 69.88 | 85.44 (7) | 85.78 (1) |
| LC2 | Precision | 99.90 | 99.76 | 99.97 | 99.83 | **99.99** | 98.74 | 99.58 | 99.96 (5) | 99.77 (1) |
|  | Recall | 92.70 | **98.16** | 81.57 | 90.48 | 86.50 | 27.03 | 69.81 | 96.99 (5) | 97.13 (1) |
| LC3 | Precision | 99.81 | 99.48 | **99.96** | 99.52 | 99.93 | 97.62 | 99.53 | 99.94 (7) | 99.45 (1) |
|  | Recall | 92.02 | **96.81** | 80.62 | 62.82 | 85.66 | 23.73 | 74.72 | 95.53 (7) | 95.67 (1) |
| LC4 | Precision | 99.89 | 99.74 | 99.98 | 99.87 | **99.99** | 99.58 | 99.29 | 99.62 (5) | 99.73 (1) |
|  | Recall | 92.56 | **98.28** | 81.44 | 88.17 | 87.08 | 35.99 | 78.86 | 96.68 (5) | 96.83 (1) |
| LC5 | Precision | 96.72 | 93.73 | 87.15 | 95.54 | 90.41 | **98.31** | 91.96 | 98.18 (7) | 94.48 (1) |
|  | Recall | 66.11 | 70.22 | 66.47 | 71.58 | 70.70 | 26.85 | 47.99 | 77.06 (7) | **77.47 (1)** |
| LC6 | Precision | 99.87 | 99.04 | 99.97 | 99.76 | **99.98** | 98.78 | 96.68 | 99.94 (6) | 99.09 (1) |
|  | Recall | 88.98 | **94.67** | 78.38 | 84.33 | 82.57 | 29.61 | 63.00 | 93.36 (6) | 93.55 (1) |
| LC7 | Precision | 99.93 | 98.07 | 99.93 | 99.78 | **99.98** | 98.79 | 99.36 | 99.96 (5) | 98.22 (1) |
|  | Recall | 90.57 | **96.15** | 80.09 | 92.16 | 85.34 | 38.81 | 70.20 | 95.04 (5) | 95.13 (1) |
| LC8 | Precision | 99.73 | 98.77 | 99.94 | 99.77 | **99.98** | 98.79 | 97.52 | 99.84 (6) | 99.12 (1) |
|  | Recall | 84.19 | 89.40 | 78.17 | 83.67 | 83.12 | 25.23 | 68.52 | 93.46 (6) | **93.66 (1)** |
| simHC | Precision | 97.68 | 97.78 | **99.69** | 99.38 | 97.29 | 98.72 | 98.54 | 99.64 (7) | 98.61 (1) |
|  | Recall | 92.54 | 90.81 | 87.91 | 74.83 | 50.16 | 88.99 | 89.96 | 92.27 (7) | **92.58 (1)** |
| simMC | Precision | 98.77 | 98.85 | **99.81** | 99.61 | 98.30 | 99.30 | 99.46 | **99.81 (7)** | 99.23 (1) |
|  | Recall | **95.89** | 95.06 | 93.00 | 86.02 | 56.50 | 94.10 | 93.02 | 95.69 (7) | 95.77 (1) |
| simLC | Precision | 98.61 | 98.62 | **99.79** | 99.61 | 98.24 | 99.20 | 99.43 | 99.74 (7) | 99.10 (1) |
|  | Recall | **95.36** | 94.45 | 92.42 | 84.63 | 56.18 | 93.34 | 93.06 | 95.21 (7) | 95.32 (1) |
| Buc12 | Precision | 95.26 | 93.03 | 98.04 | **98.82** | 95.86 | 98.06 | 98.27 | 98.68 (7) | 95.62 (1) |
|  | Recall | 72.82 | 75.50 | 67.48 | **80.31** | 69.80 | 24.31 | 57.17 | 78.54 (7) | 78.90 (1) |
| CParMed48 | Precision | 99.51 | 99.66 | **99.91** | 99.60 | 99.80 | 99.16 | 98.54 | 99.84 (5) | 99.62 (1) |
|  | Recall | 93.91 | **95.11** | 91.73 | 84.79 | 91.59 | 42.98 | 49.62 | 94.58 (5) | 94.67 (1) |
| Gut20 | Precision | 98.92 | 98.74 | 99.66 | 99.72 | 99.35 | 98.96 | 99.12 | **99.74 (7)** | 99.03 (1) |
|  | Recall | 84.60 | 85.97 | 81.30 | 75.17 | 76.04 | 33.92 | 77.49 | 90.11 (7) | **90.37 (1)** |
| Hous31 | Precision | 97.36 | 97.16 | 98.72 | **99.99** | 97.55 | 98.50 | 97.88 | 98.56 (7) | 98.00 (1) |
|  | Recall | 87.45 | 88.14 | 85.63 | 84.46 | 84.66 | 27.88 | 56.05 | 89.73 (7) | **89.89 (1)** |
| Hous21 | Precision | 99.19 | 99.33 | **99.84** | 96.55 | 99.40 | 98.97 | 99.32 | 99.61 (6) | 99.03 (1) |
|  | Recall | 86.88 | **89.07** | 84.38 | 74.36 | 85.36 | 36.64 | 49.99 | 88.78 (6) | 88.98 (1) |
| NYCSM20 | Precision | 99.23 | 99.07 | 99.71 | **99.92** | 99.60 | 97.68 | 98.82 | 99.78 (6) | 99.35 (1) |
|  | Recall | 85.33 | 85.35 | 80.11 | 68.28 | 69.19 | 24.04 | 59.53 | 90.02 (6) | **90.30 (1)** |
| Soi50 | Precision | 99.51 | 99.47 | 99.89 | 99.65 | 99.72 | 99.05 | 98.53 | **99.83 (7)** | 99.78 (1) |
|  | Recall | 92.86 | 93.61 | 89.28 | 85.62 | 90.58 | 44.26 | 48.74 | 93.71 (7) | **93.76 (1)** |
| simBA525 | Precision | 98.69 | 98.47 | 99.32 | **99.63** | 98.92 | 98.23 | 94.34 | 99.60 (7) | 99.00 (1) |
|  | Recall | 88.63 | 89.10 | 85.30 | 65.62 | 78.18 | 41.63 | 40.30 | 90.93 (1) | **91.32 (1)** |

### Run-time and Memory

Speed and memory requirements are often critical factors in the analysis of large-scale datasets. We benchmarked all tools on the same computational cluster, using sixteen threads to measure relative speed and memory consumption (Figure 7). Among the least memory intensive were MetaPhlAn, GOTTCHA, and PhyloSift. However, PhyloSift was slow compared to MetaPhlAn, GOTTCHA, MetaFlow, CLARK, and Kraken. BLAST and NBC were the slowest tools, taking multiple weeks to run for larger datasets. Taken together with precision, recall, and database size, these speed constraints can help guide the optimal selection of specific tools (Figure 7c).

**Figure 7.**
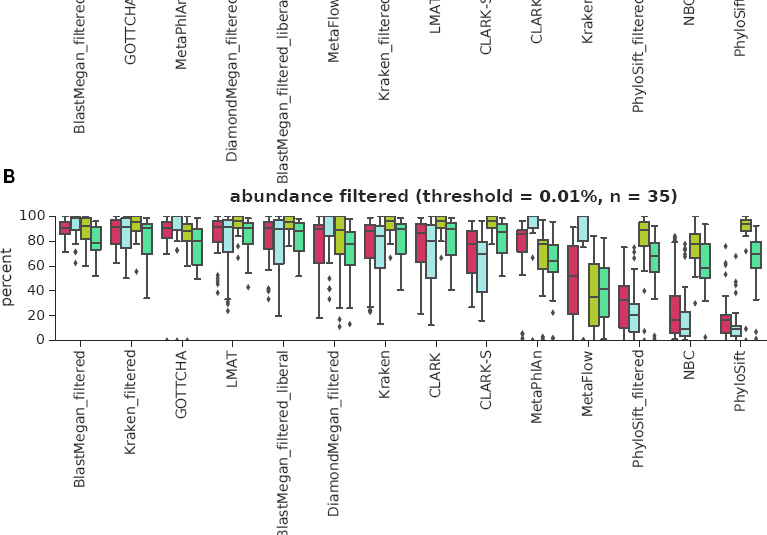
(**a**) Time and (**b**) maximum memory consumption running the tools on a subset of data using sixteen threads (except for PhyloSift, which failed to run using more than one thread). BLAST and PhyloSift were too slow to completely classify the larger datasets, therefore subsamples were taken and time multiplied. (**c**) A decision tree summary of recommendations based on the results of this analysis.

## Discussion

Recent studies of microbiomes have used a variety of molecular sequencing methods (16S, 18S, ITS, shotgun) to generate data. Many rely on a single classifier, or compare the results of a few classifiers, but classifier type and filter use differ among studies [16,47–51]. To enable greater comparability among metagenome studies, continuous benchmarking on titrated and varied datasets is needed to ensure the accuracy of these tools.

Unlike almost all prior comparisons, our analyses focused on species and strain identification, since species is a taxonomic rank more relevant in clinical diagnostics or pathogen identification than genus or phylum. Although clinical diagnosis and epidemiological tracking often require identification of strains, databases remain poorly-populated below the level of species [52]. Classification to strain requires algorithms that can differentiate genomes and their plasmids with high similarity, as we have shown for *Bacillus* [26], which is particularly challenging when using short reads. Most of the test datasets included in this study lacked complete information at the strain level, so we were able to calculate precision and recall for only a subset of datasets (n=12). These results clearly indicate that specialized approaches are still needed. For example, PanPhlAn [53] and MetaPhlAn2 strainer are recent tools designed by the authors of MetaPhlAn for epidemiological strain detection, although they focus on relationships between strains in a sample for a given species, rather than strain identification of all species in a sample. ConStrains [54] instead uses single nucleotide polymorphism profiling and requires higher depth of coverage than available for the samples used in this study.

Every database ideally should provide a complete set of all known taxa against which to compare. In reality, most species lack reference genomes, with contigs or full genomes for only around 300,000 microbial species of a recent estimate of up to 1 trillion extant species globally [55]. Large databases also demand greater computational resources, another reason that tools classify samples using limited sets of reference genomes. However, incomplete databases result in more unclassified reads, or incorrect identification of reads as related species. For this study, tools were compared using their default or recommended databases, where possible. Thus, our analyses penalize tools if their databases are missing genera or species in the truth set for a sample. We considered this a fair comparison since database size can affect the results of metagenomic analyses significantly (as we demonstrate with the limited NBC database) and certain tools were trained on, or provide, a single database.

By considering tools in their entirety, this study does not directly address differences between databases, but in the absence of any other guide for specific problems, users of these tools usually choose the default database for a given tool. Differences between tools’ default databases are shown in Table 1. For example, for full metagenomic profiling across all kingdoms of life, BLAST and Diamond offer the most extensive databases for eukaryotes, although databases can be constructed for tools like Kraken or CLARK to include greater kingdom diversity. One issue we note is that results for web-based tools that frequently update their databases (e.g., BLAST) vary over time, and may not be reproducible between analyses. The high percentage of unidentifiable reads, or “microbial dark matter”, in many studies [15,16] underscores the limitations of databases currently available, as well the use for *de novo* assembly of reads to help with the uncharacterized microorganisms from the field.

Long read technologies, such as the MinION nanopore or PacBio sequencers, can be helpful in *de novo* assembly [56,57] and avoiding ambiguous mapping of reads from conserved regions. The results suggest that even relatively low-quality reads (below an average base quality of 9) can be used for taxonomic classification, with improvements as dataset size and quality increased. Most *k*-mer-based methods performed well with longer reads, while marker-based tools did not.

## Conclusions/Best Practices

These data and results provide useful metrics, data sets (positive and negative controls), and best practices for other investigators to use, including well-characterized, titrated reference datasets now routinely sequenced by dozens of laboratories. Using the simulated datasets, read-level accuracy can be calculated and aid in determining the role of read ambiguity in taxonomic identification. Our data showed that read-level precision was much higher than organism-level precision for some tools, including Kraken, CLARK, and NBC. By varying the filtering threshold for identification and comparing F1 scores to AUPR, we showed that the discrepancy occurs because these tools detect many taxa at relatively low read counts.

To determine which taxa are actually present in a sample, users can filter their results to increase precision and exercise caution in reporting detection of low abundance species, which can be problematic to call. For example, an analysis of environmental samples collected in the Boston subway system filtered out organisms present at less than 0.1% of total abundance and in fewer than two samples [58]. Yet, depending on tool selection, this filter would have been insufficient to reject *Bacillus anthracis* in the NYC subway study, despite the absence pathogenic plasmids that distinguish it from closely-related species [26]. Therefore, filters must be considered in the context of a given study along with orthogonal information like plasmids, genome coverage, markers’ genetic variants, presence of related species, and epidemiology. Filters should be used with consideration for study design and read depth, as well as the classification tool used. However, discarding all taxa at low abundance risks rejecting species that are actually present. For instance, highly complex microbial communities found in the adult human gut and in soil contain species numbering in the hundreds and tens of thousands, respectively [59,60]. Assuming even abundance and depth of coverage, any one species would be represented by less than 0.1% of reads. In a real community of variable species abundance, many species would compose an even smaller percentage [49].

There are several options to address the ongoing problem of thresholds and low abundance species. First, precision-recall curves using known samples (such as those used in this study) can help define the appropriate filtering threshold for a given tool. Second, combining predictions from several tools offers an alternative means to improve species detection, and multiple ensemble approaches were explored in this study. Finally, targeted methods can confirm the presence of rare taxa or specific pathogens. As citizen science expands with cheaper and more accessible sequencing technologies [61,62], it is important that classifier results are not oversold.

Although many approaches are possible, here we explored ensemble methods without taking into account the differences in performance of their component tools to avoid overfitting weighted schemes. Trained predictors merit further research, including variations on that recently proposed by Metwally et al. (2016). Any ensemble method requires combining outputs of various tools, a challenge that would benefit by the adoption of standardized file formats. The Critical Assessment of Metagenomic Interpretation challenge proposed one such unifying format [26]. Inclusion of NCBI taxonomy IDs in addition to taxa names, which are more variable and difficult to track across database updates, would greatly simplify comparisons.

With significant variation in the performances of the tools demonstrated in this study, continual benchmarking using the latest sequencing methods and chemistries is critical. Tool parameters, databases, and test dataset features all affect the measures used for the comparisons. Benchmarking studies need to be computationally reproducible and transparent, and use readily available samples and methods. We showed here that filtering and combining tools decreases false positives, but that a range of issues still affect the classification of environmental samples, including depth of sequencing, sample complexity, and sequencing contamination. Additional benchmarking is necessary for analyses such as antibiotic resistance marker identification and functional classification as metagenomics moves towards answering fundamental questions beyond species composition of samples. Metrics of tool performance can inform the implementation of tools across metagenomics research studies, citizen science, and “precision metagenomics” in clinical care.

## Methods

### Data selection

A wide range of datasets was selected to answer a variety of questions. Published datasets with known species compositions (“truth sets”, see **Supplementary Table 1**) were chosen to measure precision and recall. Additional datasets with known abundances, including a subset with even (HC datasets) and log normal (LC datasets) distributions of species, facilitated analysis of abundance predictions and limits of detection. The MGRG libraries sequenced using Illumina and the MinION nanopore sequencer contain equimolar concentrations of DNA from five organisms.

We used two sets of negative controls: biological controls to test for contamination during sample preparation, and a simulated set of reads that did not map to any known organisms to test for spurious predictions. The biological control was made by spiking human NA12878 samples into a MoBio PowerSoil kit and then extracting and sequencing the DNA in triplicate. The three simulated negative control datasets we use include 100 base pair reads constructed from 17-mers that do not map to any genomes in the full NCBI/RefSeq database [36].

Lack of agreement in read classification among the tools, which can arise from discrepancies in the databases, classification algorithms, and underlying read ambiguity, was investigated. Notably, 100-base pair reads are short enough that some will map to several distinct organisms (e.g., from the same genus) within a given error rate. To facilitate a comparison between tools based solely on the database of the tool and internal sequence analysis algorithm, datasets of reads that map unambiguously to a single species within the NCBI/RefSeq database were used [36]. These six published datasets were created using the ART simulator with default error and quality base profiles [63] to simulate 100 bp Illumina reads from sets of reference sequences at a coverage of 30X as previously described [36]. Each of these unambiguous datasets (“Buc12”, “CParMed48”, “Gut20”, “Hou31”, “Hou21” and “Soi50”) represents a distinct microbial habitat based on studies that characterized real metagenomes found in the human body (mouth, gut, etc.) and in the natural or built environment (city parks/medians, houses, and soil), while a seventh dataset “simBA-525” comprised 525 randomly selected species. An extra unambiguous dataset “NYCSM20” was created to represent the organisms of the New York City subway system as described in the study of Afshinnekoo *et al.* (2015), using the same methodology as in [36]. Together, these eight unambiguous datasets contain a total of 657 species.

In the survey of the New York City subway metagenome Afshinnekoo *et al.,* noted that two samples (P00134 and P00497) showed reads that mapped to *Bacillus anthracis* using MetaPhlAn2, SURPI, and MegaBLAST-MEGAN. We used the same datasets to test for the detection of a pathogenic false positive using the wider array of tools included in this study.

### Tool Commands

#### CLARK series

We ran CLARK and CLARK-*S* CLARK is up to two orders of magnitude faster than CLARK-*S* but the latter is capable of assigning more reads with higher accuracy at the phylum/genus-level [64] and species-level [36]. Both were run using databases built from the NCBI/RefSeq bacterial, archaeal, and viral genomes.

CLARK was run on a single node using the following commands:

$ ./set_target.sh <DIR> bacteria viruses *(to set the databases at the species level)*

$ ./classify_metagenome.sh −O <file>.fasta −R <result> *(to run the classification on the file named <file>.fasta given the database defined earlier)*

$ ./estimate_abundance −D <DIR> −F result.csv > result.report.txt *(to get the abundance estimation report)*

CLARK-*S* was run on 16 nodes using the following commands:

$ ./set_target.sh <DIR> bacteria viruses

$ ./buildSpacedDB.sh *(to build the database of spaced 31-mers, using 3 different seeds)*

$ ./classify_metagenome.sh −O <file> −R <result> −n 16 ‐‐spaced

$ ./estimate_abundance −D <DIR> −F result.csv −c 0.75 −g 0.08 > result.report.txt

For CLARK-*S*, distribution plots of assignments per confidence or gamma score show an inconsistent peak localized around low values likely due to sequencing errors or noise, which suggests 1-3% of assignments are random or lack sufficient evidence. The final abundance report was therefore filtered for confidence scores greater or equal to 0.75 (“-c 0.75”) and gamma scores greater or equal to 0.08 (“-g 0.08”).

We note that we used parameters to generate classifications to the level of species for all analyses, although classifying only to genus could improve results at that level.

Speed measurements were extracted from the log.out files produced for each run.

#### GOTTCHA

Since GOTTCHA does not accept input in fasta format, fasta files for simulated datasets were converted to fastqs by setting all base quality scores to the maximum.

The v20150825 bacterial databases (GOTTCHA_BACTERIA_c4937_k24_u30_xHUMAN3x.strain.tar.gz for the strain-level analyses and GOTTCHA_BACTERIA_c4937_k24_u30_xHUMAN3x.species.tar.gz for all others) were then downloaded and unpacked and GOTTCHA run using the command: $ gottcha.pl ‐‐threads 16 ‐‐outdir $TMPDIR/ ‐‐input $TMPDIR/$DATASET.fastq −database $DATABASE_LOCATION

As for CLARK and CLARK-S, using the genus databases for classifications to genus could improve results at that level (although we observed only small differences in our comparisons to use of the species databases for a few datasets).

#### Kraken

Genomes were downloaded and a database built using the following commands:

$ kraken-build ‐‐download-taxonomy ‐‐db KrakenDB

$ kraken-build ‐‐download-library bacteria ‐‐db KrakenDB

$ kraken-build ‐‐build ‐‐db KrakenDB ‐‐threads 30

$ clean_db.sh KrakenDB

The database was then loaded into RAM:

$ sudo sync; echo 3 > /proc/sys/vm/drop_caches # clear memory buffer

$ sudo mkdir /ramdisk

$ sudo mount −t ramfs none /ramdisk

$ sudo chmod a+rwx /ramdisk

$ cp −A KrakenDB /ramdisk

Finally, Kraken was run on fasta and fastq input files using 30 nodes and a RAMDisk

$ time kraken ‐‐db /ramdisk/KrakenDB ‐‐threads 30 ‐‐fast[a/q]-input [input file] > [unfiltered output]

Results were filtered using a threshold of 0.2, which had been shown to provide a precision of ~99.1 and sensitivity ~72.8 ().

$ time kraken-filter ‐‐db /ramdisk/KrakenDB ‐‐threshold 0.2 [unfiltered output] > [filtered output]

Both filtered and unfiltered reports were generated using

$ kraken-report ‐‐db /ramdisk/KrakenDB [filtered/unfiltered output] > [report]

#### LMAT

We used the larger of the available databases, lmat-4-14.20mer.db, with the command

$ run_rl.sh ‐‐db_file=/dimmap/lmat-4-14.20mer.db ‐‐query_file=$file ‐‐threads=96 ‐‐odir=$dir ‐‐overwrite

#### MEGAN

• *BLAST*

We downloaded the NCBI BLAST executable (v2.2.28) and NT database (nucleotide) from ftp://ftp.ncbi.nlm.nih.gov/blast/. We searched for each unpaired read in the NT database using the Megablast mode of operation and an e-value threshold of 1e-20. The following command appended taxonomy columns to the standard tabular output format:

> $ blastn −query <sample>.fasta −task megablast −db NT −evalue 1e-20 \
>
> -outfmt ‘6 std staxids scomnames sscinames sskingdoms’” \
>
> > <sample>.blast

We downloaded and ran MEGAN (v5.10.6) from http://ab.inf.uni-tuebingen.de/software/megan5/. We ran MEGAN in non-interactive (command line) mode as follows:

> $ MEGAN/tools/blast2lca ‐‐format BlastTAB −topPercent 10 \
>
> ‐‐input <sample>.blast ‐‐output <sample>_read_assignments.txt

This MEGAN command returns the lowest common ancestor (LCA) taxon in the NCBI Taxonomy for each read. The topPercent option (default value 10) discards any hit with a bitscore less than ten percent of the best hit for that read.

We used a custom Ruby script (provided), summarize_megan_taxonomy_file.rb, to sum the per-read assignments into cumulative sums for each taxon. The script enforced the MEGAN parameter, Min Support Percent = 0.1, which requires that at least this many reads (as a percent of the total reads with hits) be assigned to a taxon for it to be reported. Taxa with fewer reads are assigned to the parent in the hierarchy. Output files were given the suffix “BlastMeganFiltered” to indicate that an abundance threshold (also called a filter in this manuscript) was applied. We produced a second set of output files using 0.01 as the minimum percentage and named with the suffix “BlastMeganFilteredLiberal.”

• *DIAMOND*

DIAMOND (v0.7.9.58) was run using the nr database downloaded on 2015-11-20 from NCBI (ftp://ftp.ncbi.nih.gov/blast/db/FASTA/). We tried both normal and ‐‐sensitive mode, with very similar results and present the results for the normal mode. The command to execute DIAMOND with input file sample_name.fasta is as follows, and generates an output file named sample_name.daa

diamond blastx −d /path/to/NCBI_nr/nr −q sample_name.fasta −a sample_name −p 16

MEGAN (v5.10.6) (obtained as described above) was used for read-level taxonomic classification in non-interactive mode: megan/tools/blast2lca ‐‐input sample_name.daa ‐‐format BlastTAB ‐‐topPercent 10 ‐‐gi2taxa megan/GI_Tax_mapping/gi_taxid-March2015X.bin ‐‐output sample_name.read_assignments.txt

A custom Ruby script (provided in supplement and described above) was used to sum the per-read assignments into cumulative sums for each taxon.

#### MetaFlow

MetaFlow is an alignment-based program using BLAST for fasta files produced by Illumina or 454 pyrosequencing (all fastqs for this study were converted to fastas to run MetaFlow). Any biological sample that was not sequenced with one of these technologies was not run or analyzed by MetaFlow. We ran MetaFlow using the recommended parameters as described in the available tutorial (https://github.com/alexandrutomescu/metaflow/blob/master/TUTORIAL.md). We first installed the default microbial database from NBCI/RefSeq and built the associated BLAST database. Using the provided script “Create_Blast_DB.py”, the genomes are downloaded and stored in the directory “NCBI” in the working directory and the BLAST database is created with the command: .

$ makeblastdb −in NCBI_DB/BLAST_DB.fasta −out NCBI_DB/BLAST_DB.fasta – dbtype nucl

Classification of each sample (<sample>.fasta) then proceeded through the following steps:

1) BLAST alignment

$ blastn −query <sampleID>.fasta −out <sampleID>.blast −outfmt 6 −db NCBI_DB/BLAST_DB.fasta −num_threads 10

We converted the sample file into FASTA file if the sample file was in FASTQ format and used the default settings to align the reads with BLAST.

2) LGF file construction

$ python BLAST_TO_LGF.py <sampleID>.blast NCBI_DB/NCBI_Ref_Genome.txt <avg_length> <seq_type>

The graph-based representation from the BLAST alignments is built into a LGF (Lemon Graph Format) file. This operation takes as input the average length (<avg_length>) of the reads and the sequencing machine (<seq_type>, 0 for Illumina and 1 for 454 pyrosequencing).

3) MetaFlow

$ ./metaflow −m <sampleID>.blast.lgf −g NCBI_DB/NCBI_Ref_Genome.txt −c metaflow.config

The MetaFlow program is finally run using as input the LGF file (from the previous step), the database metadata (i.e., genome length) and a configuration file. We used the default settings for the configuration but lowered the minimum threshold for abundance to increase the number of detected organisms from 0.3 to 0.001). The program outputs all the detected organisms with their related abundance and relative abundance.

#### MetaPhlAn2

MetaPhlAn2 was run using suggested command under “Basic usage” with the provided database (v20) and the latest version of bowtie2 (bowtie2-2.2.6):

$ metaphlan2.py metagenome.fasta ‐‐mpa_pkl ${mpa_dir}/db_v20/mpa_v20_m200.pkl – −bowtie2db ${mpa_dir}/db_v20/mpa_v20_m200 ‐‐input_type fasta > profiled_metagenome.txt

#### NBC

All datasets were analyzed through the web interface using the original bacterial databases [42], but not the fungal/viral or other databases [65].

Results were further filtered for the read-level analysis because every read is classified by default, using a threshold=-23.7*Read_length+490 (suggested by http://nbc.ece.drexel.edu/FAQ.php).

#### PhyloSift

PhyloSift was run using

$ phylosift all [‐‐paired] <fasta or fastq>.gz

Results were filtered for assignments with > 90% confidence.

### Analysis

#### Taxonomy IDs

For those tools that do not provide taxonomy IDs, taxa names were converted using the best matches to NCBI names before comparison of results to other tools and truth sets. A conversion table is provided in the supplementary materials (**Supplementary File 2**).

#### Precision-recall

Precision was calculated as 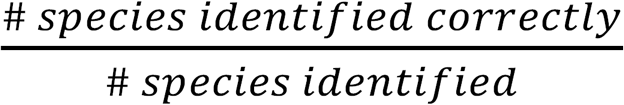 and recall as 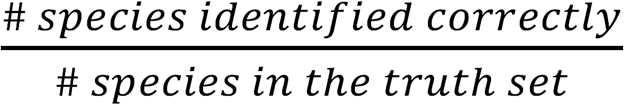. We calculated precision-recall curves by successively filtering out results based on abundances to increase precision and recalculating recall at each step, defining true and false positives in terms of the binary detection of species. The area under the precision recall curve (AUPR) was calculated using the lower trapezoid method [66]. For subspecies, classification at varying levels complicated the analysis (e.g., *Salmonella enterica* subsp. enterica, *Salmonella enterica* subsp. enterica serovar Typhimurium, *Salmonella enterica* subsp. enterica serovar Typhimurium str. LT2). We accorded partial credit if higher levels of subspecies classification were correct but the lowest were not by expanding the truth sets to include all intermediate nodes below species.

#### Negative binomial model

Negative binomial regression was used to estimate the contributions of dataset features to the number of false postives called by each tool. Using all forty datasets, false positive rate was modeled as false positives ~ ß0+ß1(X1)+ß2(X2)+ß3(X3)+ß4(X4), where X=[number of reads, number of taxa, read length, and a binary variable indicating whether a dataset is simulated]. T-statistics and associated p-values were calculated for each variable.

#### Abundance

Abundances were compared to truth set values for simulated and laboratory-sequenced data. Separate truth sets were prepared for comparison to tools that do and do not provide relative abundances by scaling expected relative abundances by genome size and ploidy (expected read proportion = (expected relative abundance)/(genome length*ploidy)), or comparing directly to read proportions. The genome size and ploidy information were obtained from the manual for the BioPool™ Microbial Community DNA Standard, while the read proportions for the HC and LC samples were calculated using species information from the fasta file headers. The log-modulus was calculated as y′ = sign(y)*log10(1+|y|) to preserve the sign of the difference between estimated and expected abundance, y.

#### Community/ensemble predictors

Ensemble predictors were designed to incorporate the results from multiple tools using either summaries of identified taxa and/or their relative abundances, or read-level classifications.

#### Summary-based

Community:

When multiple tools agree on inferred taxa, it increases confidence in the result. Conversely, when multiple tools disagree on inferred taxa, it diminishes confidence in the result. To study this intuition quantitatively, we formulated a simple algorithm for combining the outputs from multiple tools into a single "community" output. For each tool, we first we ranked the taxa from largest to smallest relative abundance, such that the most abundant taxon is rank 1 and the least abundant taxon is rank n. Next, we weighted taxa by 1/rank, such that the most abundant taxon has a weight 1, and the least abundant taxon has weight 1/n. Finally, we summed the weights for each taxon across the tools to give the total community weight for each taxon. For example, if *E. coli* were ranked 2 in five of five tools, the total weight of E. coli would be 5/2. Variations on this method of combining multiple ranked lists into a single list have been shown to effectively mitigate the uncertainty about which tool(s) are the most accurate on a particular data set [67,68].

Quorum:

As an alternative approach, we tested various combinations of three to five classifiers to predict taxa present based on the majority vote of the ensemble (known as majority-vote ensemble classifiers in machine learning literature). In the end, tools with the highest precision/recall (BlastMEGAN_Filtered, Gottcha, DiamondMEGAN_Filtered, Metaphlan, Kraken_Filtered, and LMAT) were combined to yield the best majority vote combinations. We limited the ensembles to a maximum of five classifiers, reasoning that any performance gains with more classifiers would not be worth the added computational time. Two majority vote combinations were chosen: a) BlastEnsemble, a majority vote classifier that relies on one of the BLAST-based configurations, with a taxa being called if 2 or more of the classifiers call it out of the calls from BlastMEGAN (filtered), GOTTCHA, LMAT, and MetaPhlAn, and b) DiamondEnsemble, a majority vote classifier that does not rely on BLAST, with three or more of Diamond-MEGAN, GOTTCHA, Kraken (filtered), LMAT, and MetaPhlAn calling a taxa. The second was designed to perform well but avoid BLAST-MEGAN, the tool with the highest F1 score but also the highest computational requirements.

In order to get the final relative abundance value, we tried various methods, including taking the mean or median of the ensemble. We settled on a method that prioritizes the classifiers based on the log-modulus variance of the relative abundance on the simulated data. Therefore, in the BlastEnsemble, the BLAST-MEGAN relative abundance values were taken for all taxa that were called by BLAST-MEGAN and the ensemble, then LMAT values were taken for taxa called by LMAT and the ensemble but not BLAST, then MetaPhlAn abundance values were taken for taxa called by the BlastEnsemble but not BLAST or LMAT, and finally GOTTCHA relative abundance values were used for any taxa that were called by the Blast-Ensemble but not BLAST-MEGAN, LMAT, or MetaPhlAn. This method was also apply to the DiamondEnsemble, with Kraken (filtered) prioritized, followed by LMAT, MetaPhlAn, Diamond, and GOTTCHA.

Any taxa called by any of the classifiers that were members of the ensemble but were not called by the ensemble, were set to 0. To compensate for any probability mass loss, the final relative abundance values (numerator) were divided by the sum of the relative abundance after the taxa that did not have majority vote were “zeroed-out” (denominator).

#### Read-based

For each read *r* of a given dataset, this predictor considers the classification results given by all the tools and classifies *r* using the majority vote and a “quorum” value (set in input). If all the tools agree on the assignment of *r*, say organism *o,* then the predictor classifies *r* to *o* and move to the next read, otherwise, the predictor identifies the organism *o’* of the highest vote count *v* and classifies *r* to *o’* if *v* is higher than a quorum value set by the user (ties are broken arbitrarily).

The details of the algorithm are provided in below. Parameters are the results of the tools (i.e., a list of pairs containing the read identifiers and the associated organism predicted), a quorum value (e.g., 1, 2, … 7). Note that we have set the predictor to ignore cases in which only one tool provides a prediction.

**Algorithm of the read-based ensemble predictor:**

*Input:* N lists of pairs (*read_id, taxonomy_id*) {T_1_, T_2_,…, T_N_}, a quorum Q

*Output:* A list of pairs (*read_id, taxonomy_id*)

For each *read_id r* do

 Collect all *taxonomy_id* {*s_1_,s_2_,…,s_N_} of *r* from T_1_, T_2_, …, T_N_ and store them in the list L*

 If *L* contains the same value *s* Then

  If |*L*| > 1 Then

   Output *r*, *s*

  Else

   Output *r*, “NA”

  End If

 Else // A disagreement is detected, a “board meeting” with quorum is requested:

  If *L* contains less than Q elements Then

   Output *r*, “NA”

   Continue to the next read

  End if

  Determine *s*’ the value in *L* of highest occurrence (break ties arbitrarily)

  If the occurrence of *s*’ is higher than |*L*|/2 Then

   Output *r*, *s*’

  End if

 End if

End for

### Declarations

#### Ethics approval and consent to participate

All NA12878 human data is approved for publication.

#### Consent for publication

All NA12878 human data is consented for publication.

#### Availability of data and material

The datasets supporting the conclusions of this article are freely and publicly available through the IMMSA server, http://ftp-private.ncbi.nlm.nih.gov/nist-immsa/IMMSA/.

## Funding and Acknowledgements

We would like to thank the Epigenomics Core Facility at Weill Cornell Medicine, as well as the Starr Cancer Consortium grants (I7-A765, I9-A9-071) and funding from the Irma T. Hirschl and Monique Weill-Caulier Charitable Trusts, Bert L and N Kuggie Vallee Foundation, the WorldQuant Foundation, The Pershing Square Sohn Cancer Research Alliance, NASA (NNX14AH50G, NNX17AB26G), the National Institutes of Health (R25EB020393, R01AI125416, R01ES021006), the National Science Foundation (grant #1120622), the Bill and Melinda Gates Foundation (OPP1151054), and the Alfred P. Sloan Foundation (G-2015-13964). Support was also provided by the Tri-Institutional Training Program in Computational Biology and Medicine.

## Authors’ contributions

All authors read and approved the manuscript. ABRM led the analysis. Tools were run by ABRM, RO, DD, JF, EA, RJP, GLR, and EH. ABRM, EH, GLR, SM, and PS prepared figures, with code run by multiple authors. GLR, ABRM, RJP, and RO developed the ensemble methods. ABRM, RO, EA, RRC, and CEM led the writing of the manuscript. NA, ST, and SL prepared and sequenced samples.

---

**Supplementary Figure 1.**
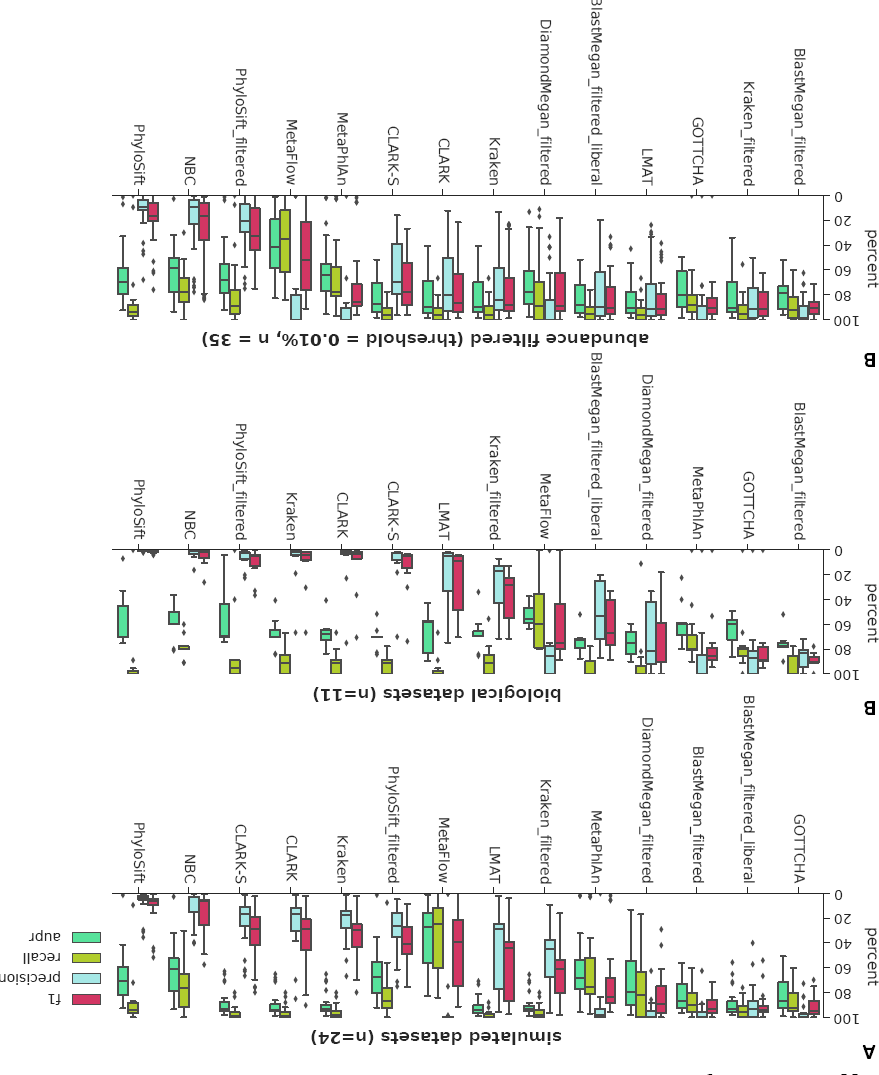
The F1-score, precision, recall, and area under the precision-recall curve for taxonomic classifications at the species level for **a)** 24 simulated datasets and **b)** 15 biological datasets, where tools are sorted by mean F1-score value.

**Supplementary Figure 2.**
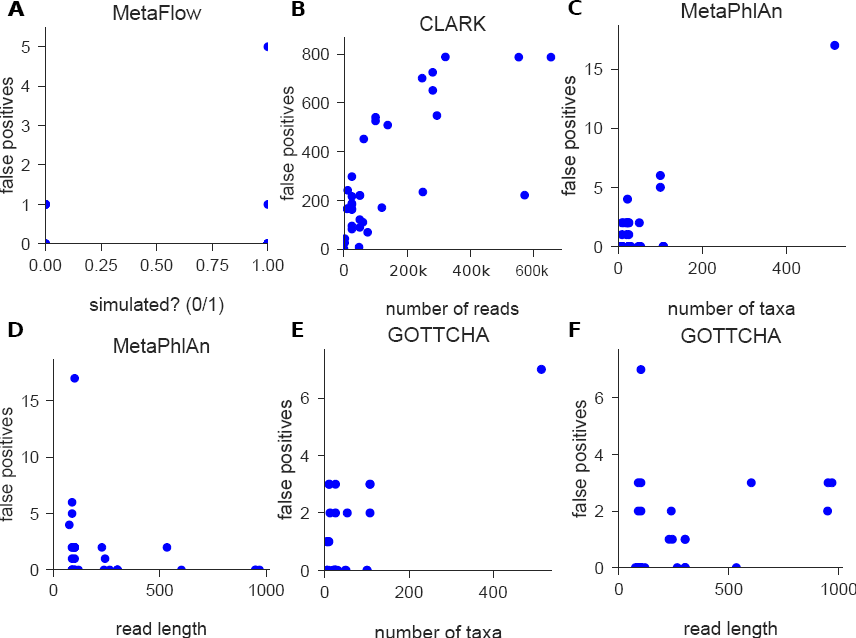
**a)** Higher false positive rates for MetaFlow decrease for only a subset of simulated datasets. **b)** False positive rates for CLARK and other *k*-mer-based classifiers increase as the number of reads in a sample increases. **c-d)** MetaPhlAn shows significant but noisy relationships between the number of taxa in a sample and read length and the number of false positives it calls. **e-f)** Similar relationships for GOTTCHA are more likely spurious.

**Supplementary Figure 3.**
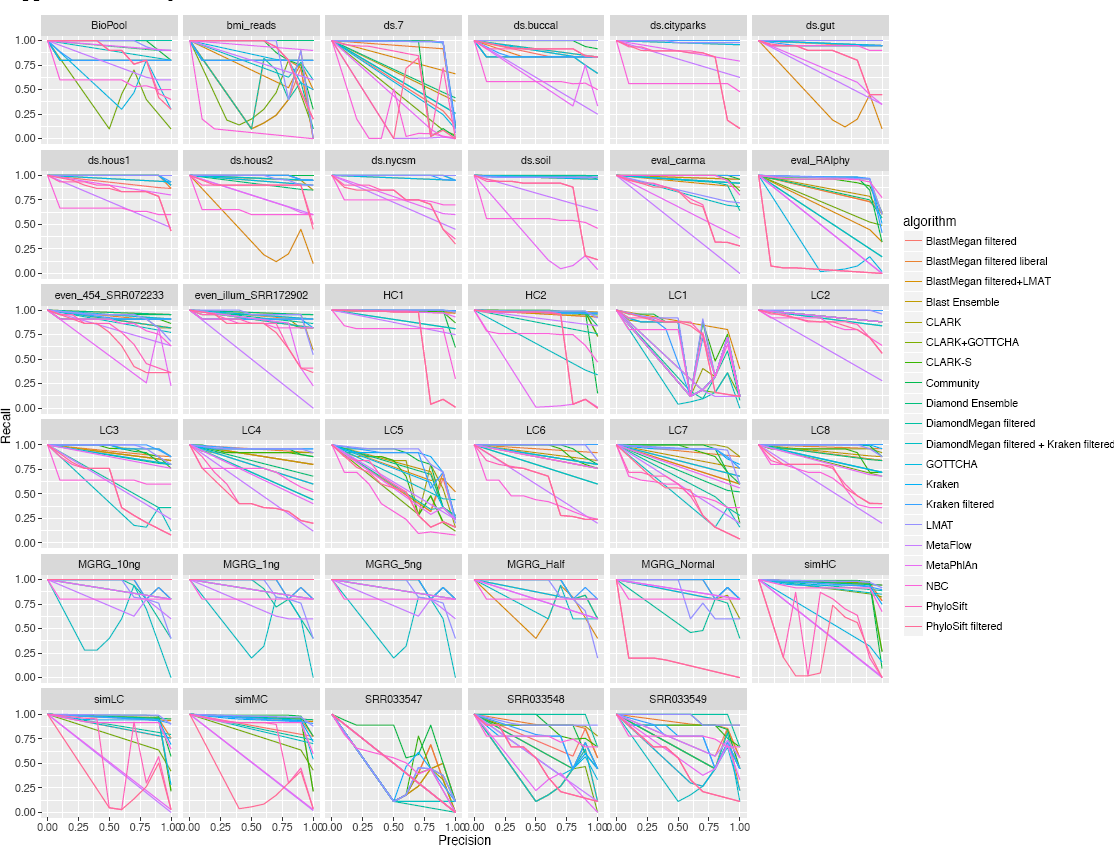
Precision-recall curves for tools on individual samples.

**Supplementary Figure 4.**
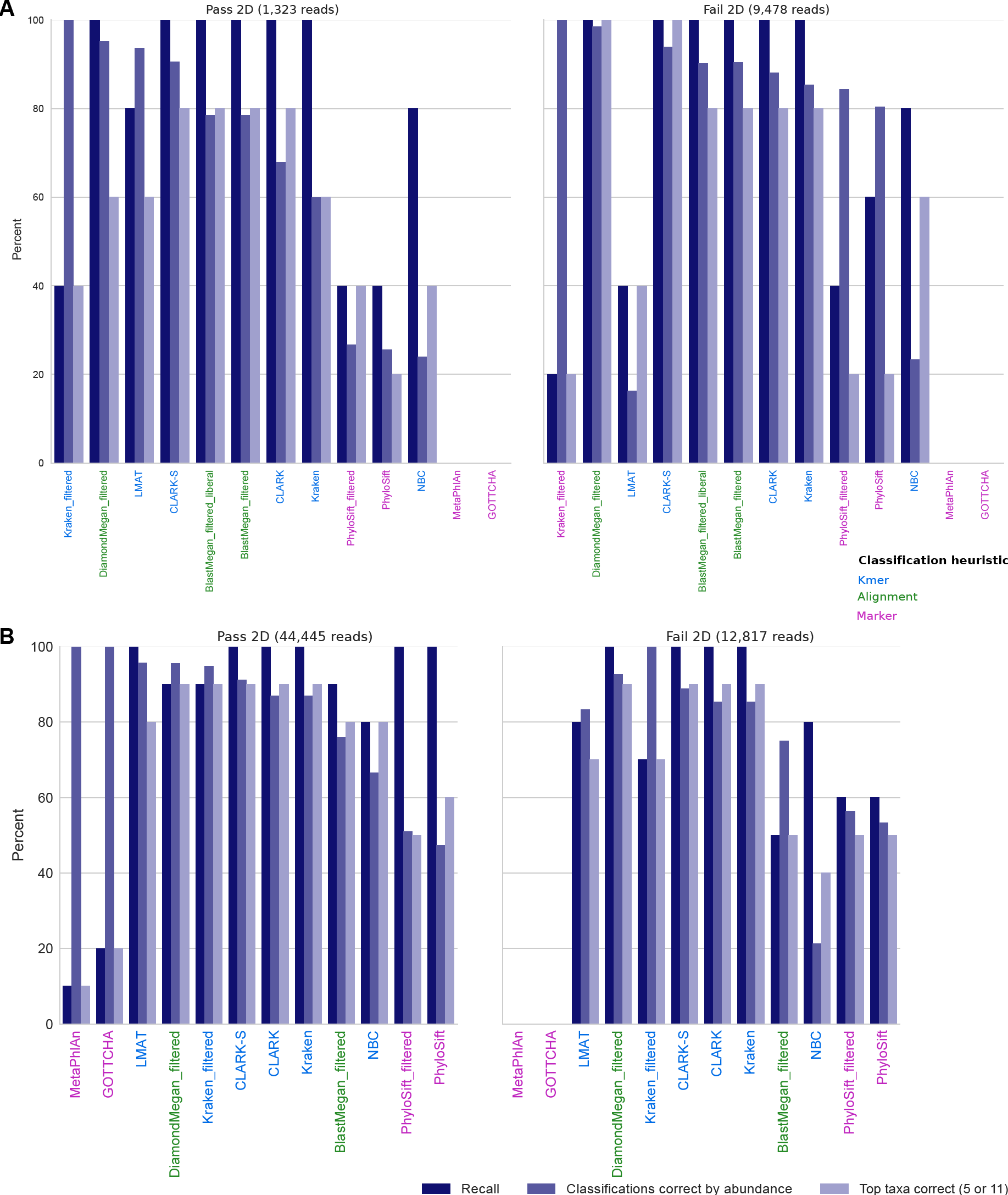
(**a**) Results from older nanopore data from 2015 for a 5-species mixture. (**b**) Results from an updated version of the technology with higher throughput and accuracy for an 11-species mixture. Kmer- and alignment-based classifiers attain high accuracy on nanopore data, even with noisy and lower quality (“Fail”, average per base quality score < 9) reads, while marker-based strategies are less effective, although this could in part be an issue of coverage. Higher coverage in (**b**) allows MetaPhlAn and GOTTCHA to correctly identify one or two species. Tools are sorted by the percent of predictions correct by abundance on the 2D pass samples.

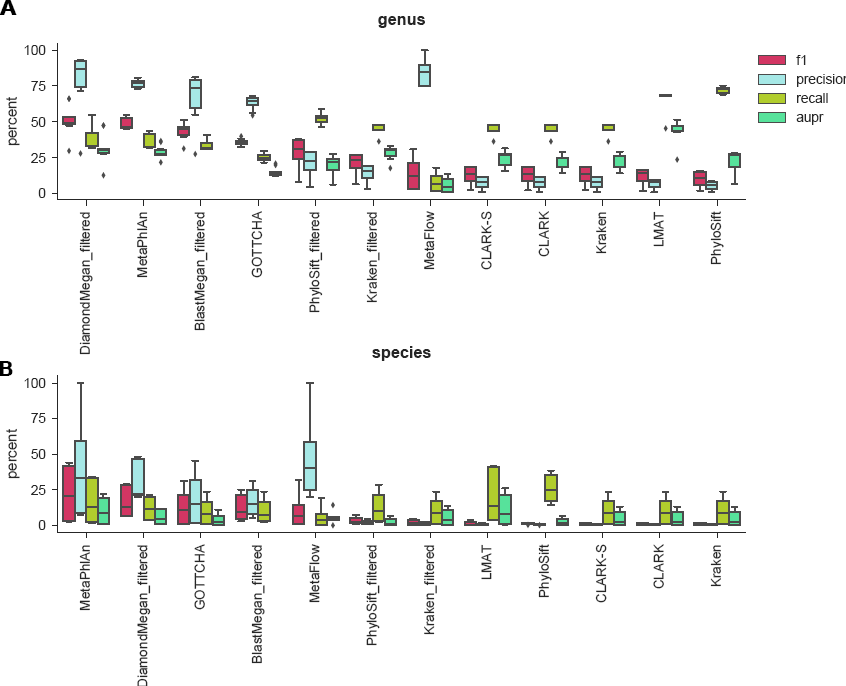

**Supplementary Table 1.** Features of datasets included in the analysis. Mean AUPR across tools provides an indication of the difficulty of a dataset.

**Supplementary Table 2.** Precision and recall at the species level for tool, listed by dataset.

**Supplementary Table 3.** Mean and median AUPR for the community predictor vs. other tools.

**Supplementry Table 4.** The read counts and relative abundances for *Bacillus anthracis* identified by various tools after the whole genome sequencing of two samples from the New York City subway system.

**Supplementary File 1.** Tool accuracy per taxon. Each file is categorized by taxonomic level. Inside each file, the first sheet shows the accuracy, the second sheet details the number of false positives, and the third sheet details the number of false negatives, of each classifier for each taxon in each taxonomic level. The three ensemble classifiers ‐‐Community, Blast Ensemble, Diamond Ensemble −are in included in this analysis for comparison.

**Supplementary File 2.** Name to taxonomy ID conversion tables for tools that do not report taxonomy IDs.

## References

1. Morgan XC, Tickle TL, Sokol H, Gevers D, Devaney KL, Ward DV, et al. Dysfunction of the intestinal microbiome in inflammatory bowel disease and treatment. Genome Biol. 2012;13:R79.

2. Tighe S, Baldwin D, Green S, Reyero N, ABRF MGRG/XMP Consortium. Next Generation Sequencing and the Extreme Microbiome Project (XMP). J. Gener. Seq. Appl. 2015;2.

3. Rose JB, Epstein PR, Lipp EK, Sherman BH, Bernard SM, Patz JA. Climate variability and change in the United States: potential impacts on water-and foodborne diseases caused by microbiologic agents. Environ. Health Perspect. 2001;109:211.

4. Verde C, Giordano D, Bellas C, di Prisco G, Anesio A. Chapter Four-Polar Marine Microorganisms and Climate Change. Adv. Microb. Physiol. 2016;69:187–215.

5. The Human Microbiome Jumpstart Reference Strains Consortium. A Catalog of Reference Genomes from the Human Microbiome. Science. 2010;328:994–9.

6. Gilbert JA, Jansson JK, Knight R. The Earth Microbiome project: successes and aspirations. BMC Biol. 2014; 12:1.

7. Weisberg WG, Barns SM, Pelletier DA, Lane DJ. 16S Ribosomal DNA Amplification for Phylogenetic Study. J. Bacteriol. 1991;173:697–703.

8. Jay ZJ, Inskeep WP. The distribution, diversity, and importance of 16S rRNA gene introns in the order Thermoproteales. Biolgy Direct. 2015;10:35.

9. Raymann K, Moeller AH, Goodman AL, Ochman H. Unexplored Archaeal Diversity in the Great Ape Gut Microbiome. Green Tringe S, editor. mSphere [Internet]. 2017;2. Available from: http://msphere.asm.org/content/2/1/e00026-17.abstract

10. Mason CE, Afshinnekoo E, Tighe S, Wu S, Levy S. International Standards for Genomes, Transcriptomes, and Metagenomes. J. Biomol. Tech. JBT. 2017;28:8–18.

11. Lan Y, Rosen G, Hershberg R. Marker genes that are less conserved in their sequences are useful for predicting genome-wide similarity levels between closely related prokaryotic strains. Microbiome. 2016;4:1–13.

12. Lindgreen S, Adair KL, Gardner PP. An evaluation of the accuracy and speed of metagenome analysis tools. Sci. Rep. 2016;6.

13. Ounit R, Wanamaker S, Close TJ, Lonardi S. CLARK: fast and accurate classification of metagenomic and genomic sequences using discriminative *k*-mers. BMC Genomics. 2015;16:236.

14. Muñoz-Amatriaín M, Lonardi S, Luo M, Madishetty K, Svensson JT, Moscou MJ, et al. Sequencing of 15 622 gene-bearing BACs clarifies the gene-dense regions of the barley genome. Plant J. 2015;84:216–27.

15. Yooseph S, Andrews-Pfannkoch C, Tenney A, McQuaid J, Williamson S, Thiagarajan M, et al. A metagenomic framework for the study of airborne microbial communities. PLoS One. 2013;8:e81862.

16. Afshinnekoo E, Meydan C, Chowdhury S, Jaroudi D, Boyer C, Bernstein N, et al. Gesospatial Resolution of Human and Bacterial Diversity from City-scale Metagenomics. Cell Syst. 2015;1:72–87.

17. Petit RA, Ezewudo M, Joseph SJ, Read TD. Searching for anthrax in the New York City subway metagenome. [Internet]. 2015 [cited 2017 Jan 9]. Available from: https://read-lab-confederation.github.io/nyc-subway-anthrax-study/

18. Ackelsberg J, Rakeman J, Hughes S, Petersen J, Mead P, Schriefer M, et al. Lack of Evidence for Plague or Anthrax on the New York City Subway. Cell Syst. 2015;1:4–5.

19. Minot SS, Greenfield N, Afshinnekoo E, Mason CE. Detection of Bacillus anthracis using a targeted gene panel [Internet]. 2015 [cited 2016 Dec 29]. Available from: https://science.onecodex.com/bacillus-anthracis-panel/

20. Peabody MA, Van Rossum T, Lo R, Brinkman FS. Evaluation of shotgun metagenomics sequence classification methods using in silico and in vitro simulated communities. BMC Bioinformatics. 2015;16:1.

21. Gonzalez A, Vázquez-Baeza Y, Pettengill J, Ottesen A, McDonald D, Knight R. Avoiding Pandemic Fears in the Subway and Conquering the Platypus. mSystems. 2016;1:e00050–16.

22. Bradley P, Gordon NC, Walker TM, Dunn L, Heys S, Huang B, et al. Rapid antibiotic-resistance predictions from genome sequence data for Staphylococcus aureus and Mycobacterium tuberculosis. Nat. Commun. 2015;6.

23. Sinha R, Abnet CC, White O, Knight R, Huttenhower C. The microbiome quality control project: baseline study design and future directions. Genome Biol. 2015; 16:1.

24. IMMSA Mission Statement | NIST [Internet]. 2016 [cited 2017 Jan 17]. Available from: https://www.nist.gov/mml/bbd/immsa-mission-statement

25. MetaSUB International Consortium. The Metagenomics and Metadesign of the Subways and Urban Biomes (MetaSUB) International Consortium inaugural meeting report. Microbiome. 2016;4:1–14.

26. CAMI – Critical Assessment of Metagenomic Interpretation [Internet]. [cited 2016 Feb 10]. Available from: http://www.cami-challenge.org

27. Sczyrba A, Hofmann P, Belmann P, Koslicki D, Janssen S, Droege J, et al. Critical Assessment of Metagenome Interpretation-a benchmark of computational metagenomics software. bioRxiv. 2017;99127.

28. Richardson RT, Bengtsson-Palme J, Johnson RM. Evaluating and optimizing the performance of software commonly used for the taxonomic classification of DNA metabarcoding sequence data. Mol. Ecol. Resour. 2016;n/a-n/a.

29. Bazinet AL, Cummings MP. A comparative evaluation of sequence classification programs. BMC Bioinformatics. 2012; 13:1.

30. Lu J, Breitwieser FP, Thielen P, Salzberg SL. Bracken: Estimating species abundance in metagenomics data. bioRxiv. 2016;51813.

31. Parisot N. Détermination de sondes oligonucléotidiques pour l’exploration á haut débit de la diversité taxonomique et fonctionnelle d’environnements complexes. 2014;

32. Altschul SF, Madden TL, Schaffer AA, Zhang J, Zhang Z, Miller W, et al. Gapped BLAST and PSI-BLAST: a new generation of protein database search programs. Nucleic Acids Res. 1997;25:3389–402.

33. Freitas TAK, Li P-E, Scholz MB, Chain PS. Accurate read-based metagenome characterization using a hierarchical suite of unique signatures. Nucleic Acids Res. 2015;gkv180.

34. Huson DH, Mitra S, Ruscheweyh H-J, Weber N, Schuster SC. Integrative analysis of environmental sequences using MEGAN4. Genome Res. 2011;21:1552–60.

35. Huson DH, Auch AF, Qi J, Schuster SC. MEGAN analysis of metagenomic data. Genome Res. 2007;17:377–86.

36. Ounit R, Lonardi S. Higher classification sensitivity of short metagenomic reads with CLARK-S Bioinformatics. 2016;32:3823–5.

37. Buchfink B, Xie C, Huson DH. Fast and sensitive protein alignment using DIAMOND. Nat. Methods. 2015;12:59–60.

38. Wood DE, Salzberg SL. Kraken: ultrafast metagenomic sequence classification using exact alignments. Genome Biol. 2014;15:R46.

39. Ames SK, Hysom DA, Gardner SN, Lloyd GS, Gokhale MB, Allen JE. Scalable metagenomic taxonomy classification using a reference genome database. Bioinformatics. 2013;29:2253–60.

40. Sobih A, Tomescu AI, Mäkinen V. MetaFlow: Metagenomic profiling based on whole-genome coverage analysis with min-cost flows. Springer; 2016. p. 111–21.

41. Truong DT, Franzosa EA, Tickle TL, Scholz M, Weingart G, Pasolli E, et al. MetaPhlAn2 for enhanced metagenomic taxonomic profiling. Nat. Methods. 2015;12:902–3.

42. Rosen GL, Reichenberger ER, Rosenfeld AM. NBC: the Naive Bayes Classification tool webserver for taxonomic classification of metagenomic reads. Bioinformatics. 2011;27:127–9.

43. Darling AE, Jospin G, Lowe E, Matsen FA, Bik HM, Eisen JA. PhyloSift: phylogenetic analysis of genomes and metagenomes. PeerJ [Internet]. 2014;2. Available from: http://dx.doi.org/10.7717/peerj.243

44. Salter SJ, Cox MJ, Turek EM, Calus ST, Cookson WO, Moffatt MF, et al. Reagent and laboratory contamination can critically impact sequence-based microbiome analyses. BMC Biol. 2014;12:87.

45. Schneiker S, Perlova O, Kaiser O, Gerth K, Alici A, Altmeyer MO, et al. Complete genome sequence of the myxobacterium Sorangium cellulosum. Nat Biotech. 2007;25:1281–9.

46. Koslicki D, Foucart S, Rosen G. Quikr: a method for rapid reconstruction of bacterial communities via compressive sensing. Bioinformatics. 2013;29:2096–102.

47. Hemme CL, Tu Q, Qin Y, Gao W, Deng Y, Nostrand JDV, et al. Comparative metagenomics reveals impact of contaminants on groundwater microbiomes. Front. Microbiol. 2015;6:1205.

48. Stolze Y, Zakrzewski M, Maus I, Eikmeyer F, Jaenicke S, Rottmann N, et al. Comparative metagenomics of biogas-producing microbial communities from production-scale biogas plants operating under wet or dry fermentation conditions. Biotechnol. Biofuels. 2015;8:14.

49. Wilson MR, Naccache SN, Samayoa E, Biagtan M, Bashir H, Yu G, et al. Actionable diagnosis of neuroleptospirosis by next-generation sequencing. N. Engl. J. Med. 2014;370:2408–17.

50. Young JC, Chehoud C, Bittinger K, Bailey A, Diamond JM, Cantu E, et al. Viral metagenomics reveal blooms of anelloviruses in the respiratory tract of lung transplant recipients. Am. J. Transplant. 2015;15:200–9.

51. Chu DM, Ma J, Prince AL, Antony KM, Seferovic MD, Aagaard KM. Maturation of the infant microbiome community structure and function across multiple body sites and in relation to mode of delivery. Nat Med [Internet]. 2017;advance online publication. Available from: http://dx.doi.org/10.1038/nm.4272

52. Dijkshoorn L, Ursing B, Ursing J. Strain, clone and species: comments on three basic concepts of bacteriology. J. Med. Microbiol. 2000;49:397–401.

53. Scholz M, Ward DV, Pasolli E, Tolio T, Zolfo M, Asnicar F, et al. Strain-level microbial epidemiology and population genomics from shotgun metagenomics. Nat. Methods. 2016;

54. Luo C, Knight R, Siljander H, Knip M, Xavier RJ, Gevers D. ConStrains identifies microbial strains in metagenomic datasets. Nat. Biotechnol. 2015;33:1045–52.

55. Locey KJ, Lennon JT. Scaling laws predict global microbial diversity. Proc. Natl. Acad. Sci. 2016;201521291.

56. Karlsson E, Lärkeryd A, Sjödin A, Forsman M, Stenberg P. Scaffolding of a bacterial genome using MinlON nanopore sequencing. Sci. Rep. 2015;5.

57. Cao MD, Nguyen SH, Ganesamoorthy D, Elliott A, Cooper M, Coin LJ. Scaffolding and Completing Genome Assemblies in Real-time with Nanopore Sequencing. bioRxiv. 2016;54783.

58. Hsu T, Joice R, Vallarino J, Abu-Ali G, Hartmann EM, Shafquat A, et al. Urban Transit System Microbial Communities Differ by Surface Type and Interaction with Humans and the Environment. mSystems. 2016;1:e00018–16.

59. Qin J, Li R, Raes J, Arumugam M, Burgdorf KS, Manichanh C. A human gut microbial gene catalogue established by metagenomic sequencing. Nature [Internet]. 2010;464. Available from: http://dx.doi.org/10.1038/nature08821

60. Roesch LF, Fulthorpe RR, Riva A, Casella G, Hadwin AK, Kent AD, et al. Pyrosequencing enumerates and contrasts soil microbial diversity. ISME J. 2007;1:283–90.

61. Erlich Y. A vision for ubiquitous sequencing. Genome Res. 2015;25:1411–6.

62. Zaaijer S, Erlich Y. Using mobile sequencers in an academic classroom. Elife. 2016;5:e14258.

63. Huang W, Li L, Myers JR, Marth GT. ART: a next-generation sequencing read simulator. Bioinformatics. 2012;28:593–4.

64. Ounit R, Lonardi S. Higher classification accuracy of short metagenomic reads by discriminative spaced k-mers. Springer; 2015. p. 286–95.

65. Rosen GL, Lim TY. NBC update: The addition of viral and fungal databases to the Naïve Bayes classification tool. BMC Res. Notes. 2012;5:1.

66. Boyd K, Eng KH, Page CD. Area under the Precision-Recall Curve: Point Estimates and Confidence Intervals. In: Blockeel H, Kersting K, Nijssen S, Železný F, editors. Mach. Learn. Knowl. Discov. Databases Eur. Conf. ECML PKDD 2013 Prague Czech Repub. Sept. 23-27 2013 Proc. Part III [Internet]. Berlin, Heidelberg: Springer Berlin Heidelberg; 2013. p. 451–66. Available from: http://dx.doi.org/10.1007/978-3-642-40994-3_29

67. Prill RJ, Marbach D, Saez-Rodriguez J, Sorger PK, Alexopoulos LG, Xue X, et al. Towards a rigorous assessment of systems biology models: the DREAM3 challenges. PloS One. 2010;5:e9202.

68. Marbach D, Costello JC, Küffner R, Vega NM, Prill RJ, Camacho DM, et al. Wisdom of crowds for robust gene network inference. Nat. Methods. 2012;9:796–804.

